# A dendritic substrate for temporal diversity of cortical inhibition

**DOI:** 10.1101/2024.07.09.602783

**Authors:** Annunziato Morabito, Yann Zerlaut, Dhanasak Dhanasobhon, Emmanuelle Berthaux, Alexandra Tzilivaki, Gael Moneron, Laurence Cathala, Panayiota Poirazi, Alberto Bacci, David DiGregorio, Joana Lourenço, Nelson Rebola

## Abstract

In the mammalian neocortex, GABAergic interneurons (INs) inhibit cortical networks in profoundly different ways. The extent to which this depends on how different INs process excitatory signals along their dendrites is poorly understood. Here, we reveal that the functional specialization of two major populations of cortical INs is determined by the unique association of different dendritic integration modes with distinct synaptic organization motifs. We found that somatostatin (SST)-INs exhibit NMDAR-dependent dendritic integration and uniform synapse density along the dendritic tree. In contrast, dendrites of parvalbumin (PV)-INs exhibit passive synaptic integration coupled with proximally enriched synaptic distributions. Theoretical analysis shows that these two dendritic configurations result in different strategies to optimize synaptic efficacy in thin dendritic structures. Yet, the two configurations lead to distinct temporal engagement of each IN during network activity. We confirmed these predictions with *in vivo* recordings of IN activity in the visual cortex of awake mice, revealing a rapid and linear recruitment of PV-INs as opposed to a long-lasting integrative activation of SST-INs. Our work reveals the existence of distinct dendritic strategies that confer distinct temporal representations for the two major classes of neocortical INs and thus dynamics of inhibition.

---

In the neocortex, fast synaptic inhibition originates from a rich diversity of inhibitory neurons (INs) that play a fundamental role in regulating cortical function by orchestrating network dynamics (*1–3*). Importantly, this IN diversity is associated with differential functional roles of specific INs within neuronal circuits (*3*, *4*). The diverse IN functions are thought to result mainly from differences in connectivity profiles, specific biophysical properties, strength and dynamics of excitatory glutamatergic synapses that recruit GABAergic INs (5–10). However, INs primarily receive synaptic inputs along their dendrites, and it remains unclear whether distinct dendritic properties across different IN populations contribute to their specific activity profiles and thus computational roles in cortical circuit function.

Recent evidence suggests that IN dendrites can exhibit nonlinear integration of synaptic activity, similarly to PNs, thus increasing the computational power of INs (for a review see (*5–8*). However, most research on dendritic operations in INs was limited to PV cells of either hippocampus (*8–12*) or cerebellum (*13*, *14*). Indeed, it has been shown that oriens-alveus (O/A)-SST INs in the hippocampus exhibit significant differences in the density of voltage-gated potassium and sodium channels as compared to PV-INs (*15*), indicating that dendritic properties might vary between different populations of INs. Yet, it is presently uncertain whether specificities in dendritic integration determine the functional properties of different populations of INs.

Among the different subtypes, PV- and SST-IN are the two main IN populations that provide inhibitory control of PN networks. They do so, however, using distinct dynamics. Fast-spiking, PV basket cells are phase-locked to fast oscillations, are highly responsive to increases in activity of excitatory inputs and regulate PN activity via perisomatic inhibition with millisecond-level precision (*5*, *16–18*). In contrast, the activation of SST-INs requires bursts from single PNs or the firing of multiple PNs. Moreover, SST-INs fire with less precision, and inhibit PN dendritic segments over longer timescales (*19–24*). These distinct features are essential for cortical network properties, including emergence of oscillatory rhythms and facilitating learning through dendritic inhibition and gating synaptic plasticity (*18*, *23*, *25–28*). However, whether differences in dendritic integration properties play a role in determining the temporal dynamics in the recruitment of PV- and SST-INs remains largely unexplored. Here, we demonstrate that PV- and SST-INs employ distinct strategies for integrating excitatory inputs on their dendrites. These strategies govern the firing dynamics of each interneuron subclass, differently affect the encoding of sensory inputs by cortical networks, and thereby define their specific functional roles within cortical networks.

## PV- and SST-INs possess distinct dendritic integration properties

To investigate synaptic integration in dendrites of neocortical INs, we performed whole-cell, current-clamp recordings of PV and SST-INs in layer 2/3 of primary somatosensory cortex in adult mice and used two-photon glutamate uncaging to mimic synapse activation (*13*, *29*); **Fig. 1A-D**). We monitored somatic voltage responses to both single synapse activation and to multiple quasi-synchronous input sequences. To assess deviations from linear synaptic integration, we compared the arithmetic sum (which represents the linear prediction) of individual photolysis-evoked excitatory postsynaptic potentials (pEPSPs) with actual responses evoked by quasi-simultaneous (0.12 ms inter-pulse interval) activation of all locations (**Fig. 1A,C**). We found an overall sublinear integration mode in PV-INs dendrites (**Fig. 1A,B**) and an overall supralinear integration in SST-INs (**Fig. 1C,D**). Such a difference was also observed for different time intervals between active inputs (**Fig. S1**). The supralinear integration observed in SST-INs was not significantly different from the values obtained in pyramidal neurons (PNs) (**Fig. 1D,E**).

**Figure 1:**
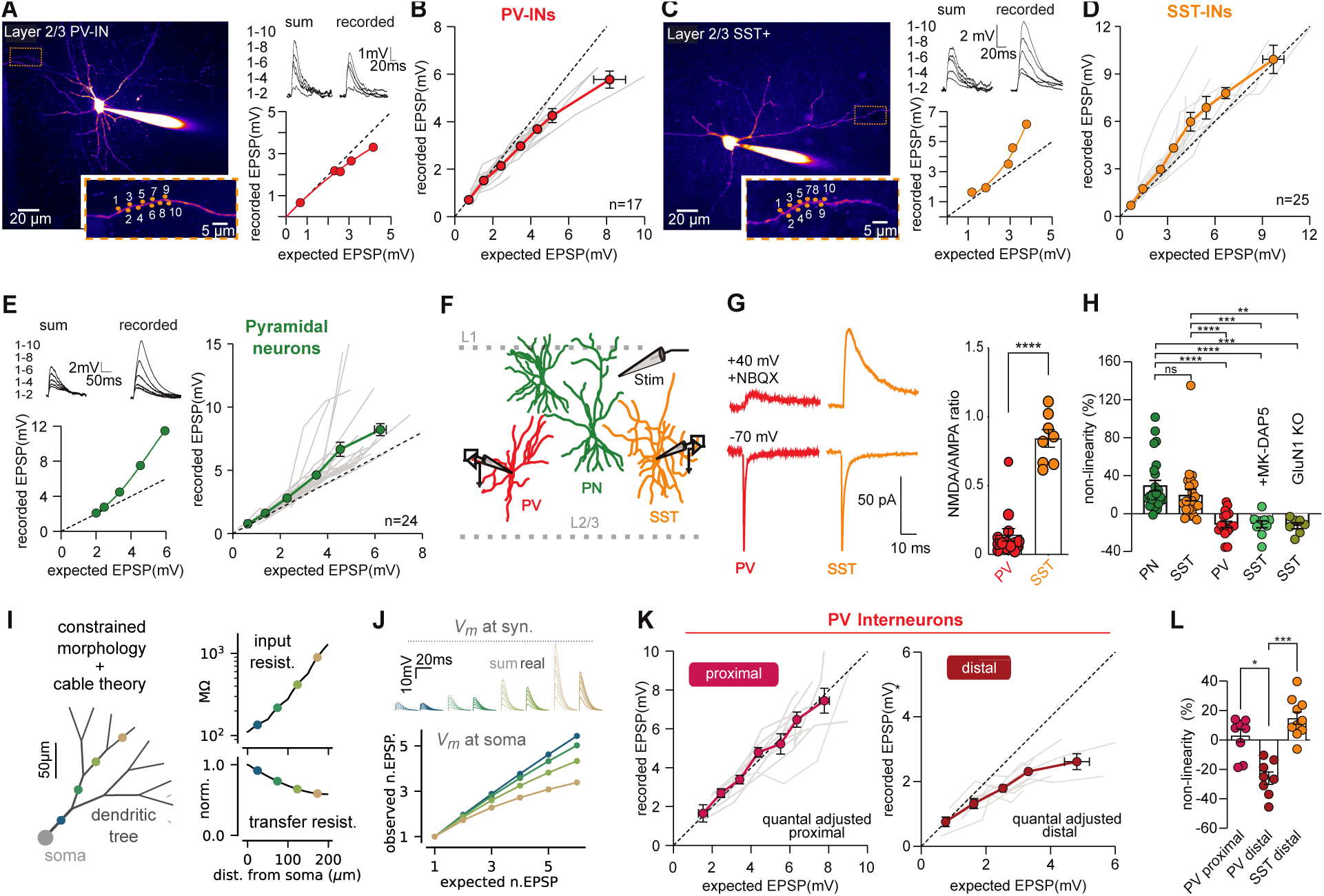
PV- and SST-INs display different dendritic integration properties. **A)** *(left)* Two-photon laser scanning microscopy (2PLSM) image (maximum-intensity projection, MIP) of a PV-IN patch loaded with Alexa-594 (10 μM). The inset shows a selected dendrite and uncaging sites for 10 synapses. (*right*) Example of photolysis-evoked EPSPs in a dendrite with increasing number of stimulated synapses (from locations in A, activated at 0.12 ms intervals). The traces compare the calculated linear sum of individual responses to the recorded response from quasi-synchronous synapse activation. The input-output curve shows recorded somatic EPSP amplitudes versus the expected linear sum. **B)** Summary plot of input-output curves for dendrites of PV-INs. Single grey lines represent individual experiments (n=17) and in red the binned average. **C-D**) Same as A-B, but for SST-INs. **E)** Same as A-B, but for pyramidal neurons. **F)** Schematic representation of experimental conditions used to stimulate local excitatory connections and record glutamatergic synaptic activity in PV- and SST-INs in layer 2/3 of primary somatosensory cortex. **G)** Representative traces *(left)* and summary plot *(right)* of EPSCs recorded at -70 mV and +40 mV in PV- and SST-INs (n=8; p<0.001, Mann-Whitney). **H)** Percentage of non-linearity obtained in PYR (green, n=25), SST-INs (orange, n=23), PV-INs (red, n=17), SST INs in the presence of NMDA blockers (light green, n=14) and SST-INs from GluN1-KO mice (khaki green, n=7). The data are presented as average ± SEM; **p<0.001,***p<0.0001, ****p<0.00001, Kruskal-Wallis test with Dunn’s correction. **I)** Properties of the simplified model of PV-IN integration. *(left)* Schematic of the morphology with a somatic compartment and symmetric-branching dendritic tree. *(right)* Input resistance and transfer resistance profiles along the dendritic tree relative to soma distance. **J)** Input-output curve in the simplified model. *(top inset)* Depolarization values from quasi-synchronous input of increasing number of synapses (1 to 6 synapses) at various dendritic locations (color-coded as in I). Dashed and solid lines show linear prediction (sum) and observed (real) responses. (bottom) Relationship between expected (sum) and observed (real) peak EPSP values at different dendritic locations (color-coded), normalized to single event EPSP values. **K)** Subthreshold input-output relationship of pEPSPs in proximal (<40 μm from soma) and distal (100 μm from soma) dendrites using quantal adjusted pEPSP amplitude. Dashed line indicates slope of 1. **L)** Summary of non-linearity percentage in proximal (cardinal red) and distal (brick red) PV-INs and distal SST-INs dendrites (orange) using quantal adjusted EPSP amplitude. Data are average ± SEM, *p<0.01,***p<0.0001, Kruskal-Wallis test with Dunn’s correction

In PNs, integration of glutamatergic synaptic inputs along individual dendritic branches has been shown to be strongly dependent on the presence of synaptic NMDARs (*30–32*). We explored whether variations in dendritic integration were associated with differences in the relative contribution of NMDA and AMPA receptors (AMPAR) at excitatory synapses onto PV- and SST-INs. Using voltage-clamp recordings and extracellular stimulation, we observed that SST-INs exhibit a higher NMDA-to-AMPA ratio when compared to PV-INs (**Fig. 1F, G**). This difference was confirmed by paired recordings from unitary synaptic connections between individual L2/3 PNs and PV- or SST-INs (**Fig. S2A,B**). Furthermore, by examining NMDAR-dependent miniature EPSCs at -30 mV (isolated using the NMDAR blocker D-AP5, 100 µM), we found that the marked NMDA/AMPA differences between PV- and SST-INs was predominantly due to a much weaker expression of the NMDAR-mediated component in PV-INs (**Fig. S2C,D**). To test whether the enhanced levels of synaptic NMDARs in SST-INs defined its supralinear integration properties, we measured subthreshold input-output function of L2/3 SST-INs in the presence of pharmacological blockade (MK-801 and D-AP5) or upon genetic removal (GluN1 KO) of NMDARs. In both conditions, supralinear integration was abolished and integration of quasi-synchronous inputs resulted in compound EPSPs smaller than the arithmetic linear sum (**Fig. 1H**).

Sublinear integration observed in PV-INs is compatible with a decrease in driving force due to large local dendritic depolarizations as observed in cerebellar stellate (*13*) and Golgi cells (*14*), as well as in hippocampal and L5 prefrontal cortex PV interneurons, where both supralinear and sublinear branches coexist (*11*, *12*). To investigate whether such an effect could explain neocortical PV-IN integration, we analyzed a reduced theoretical model implementing cable theory in a simplified morphology ((*33*), **Fig. 1I**). Model parameters were constrained to match our electrophysiological and morphological measurements (see **Methods**). This simple passive dendritic model recapitulated sublinear integration in PV INs because of high local input resistance (**Fig. 1J**, top). The model also predicted that input-output curves strongly relied on spatial location of excitatory synapses, with sublinear integration being particularly robust at distal locations (**Fig. 1J**, bottom). To test this, we performed synaptic integration experiments separately at proximal and distal locations (**Fig. 1K**, <40μm and >100μm from the soma respectively). Uncaging laser power was adjusted to mimic average EPSPs amplitude at the corresponding specific dendritic locations (**Fig. S3)**. In accordance with the model predictions, we observed that PV-INs exhibited strong sublinear responses when dendrites were stimulated distally. In contrast, dendritic integration was mostly linear when dendrites were stimulated proximally (**Fig. 1K,L**). Conversely, responses of SST-IN dendrites were supralinear, regardless of dendritic location (**Fig. 1L**). Importantly, differences in dendritic integration between PV- and SST-INs could not be attributable to differences in dendritic diameter along their length, as both cell types exhibited similar dendritic diameter profiles (**Fig. S4**). Overall, these results indicate that the dendrites of PV- and SST-INs have different dendritic integration properties. PV-INs display dendritic integration gradients along the dendritic tree with strong distal sublinear integration, compatible with passive integration in thin dendrites (*11*, *13*). On the other hand, SST-INs display NMDAR-dependent supralinear integration along the entire length of their dendrites, thus providing effective excitatory input integration despite cable filtering properties of thin dendrites.

## Synaptic integration of multivesicular release is different between PV- and SST-INs

Previous studies showed that a consequence of nonlinear integration is the distance-dependent alteration in short-term plasticity (*13*). We therefore hypothesized that the distinct dendritic integration properties could result in differential processing of short-term presynaptic plasticity by PV- and SST-IN dendrites. This effect becomes especially significant if multiple release events (*i.e.,* quantal content greater than one) occur at single points of contact and require integration in close temporal and spatial proximity (*13*). We thus examined if glutamatergic synapses innervating PV- and SST-INs elicited multiple quantal release events per individual action potential. To address this question, we used the genetically-encoded glutamate sensor, iGluSnFR (*34*) to monitor glutamate release events from single L2/3 PN axonal boutons *in vitro* (**Fig. 2A**). iGluSnFR was expressed in either SST- or PV-INs. By loading individual L2/3 PNs with Alexa 594 via the whole-cell pipette, and utilizing two-photon imaging, we were able to identify putative points of contact between presynaptic axons and iGluSnFR expressing IN dendrites (**Fig. 2A,B**). To elicit glutamate release, we evoked action potentials in presynaptic L2/3 PNs via current injection (**Fig. 2A**). We successfully identified specific functional synaptic contacts, enabling the detection of glutamate release events even within individual recording sweeps (**Fig. 2B-C**).

**Figure 2:**
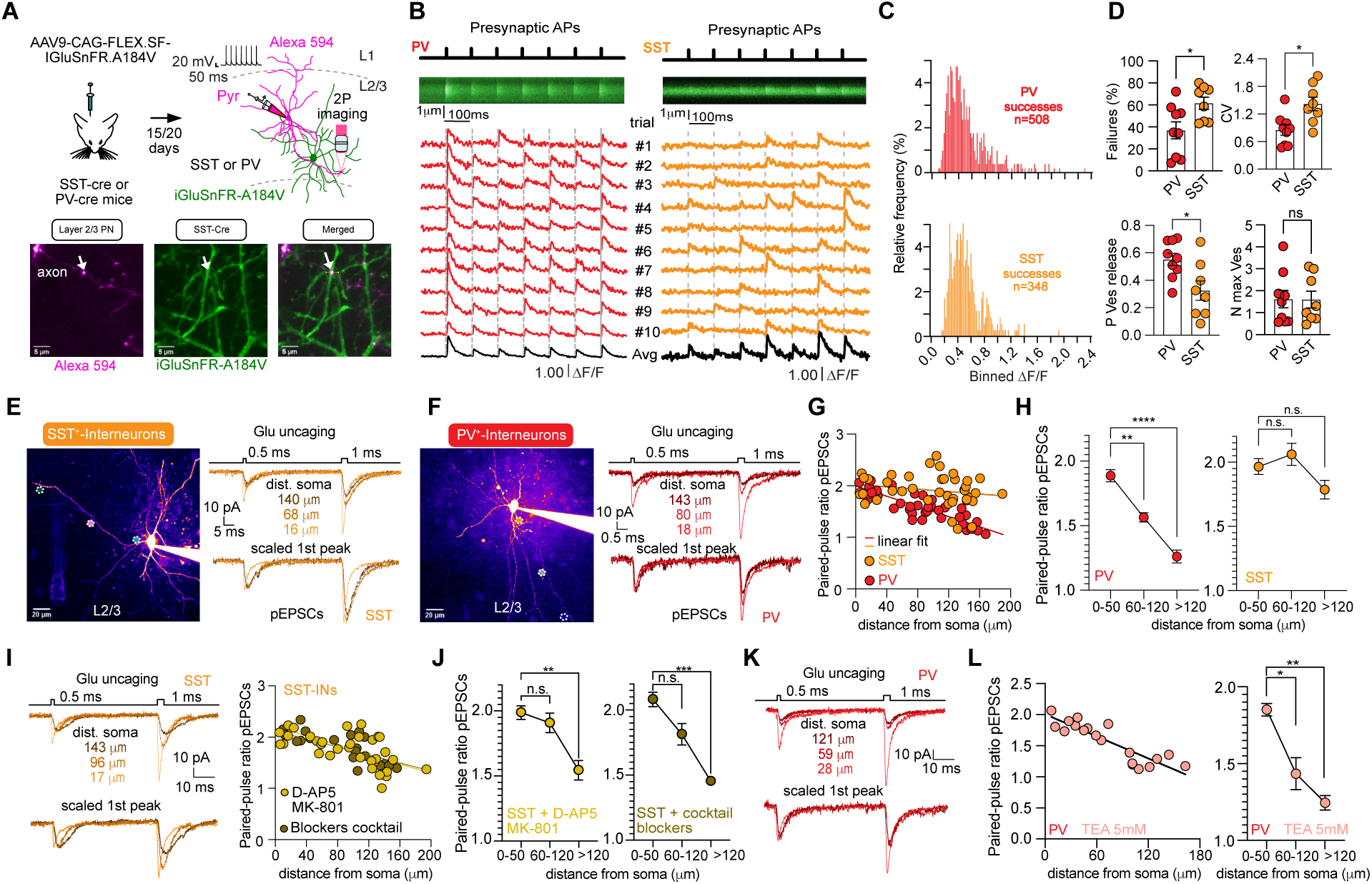
Multivesicular release summates sublinearly in PV-INs dendrites and predominantly linearly in SST-INs dendrites. **A)** (Top) Methodology to study neurotransmitter release at synaptic contacts between PNs and PV- or SST-INs using iGluSNFR-A184V expressed in PV- or SST-INs. Glutamate release was evoked from L2/3 PNs via current injection. (Bottom) Two-photon images show a patch-loaded PN axon (red) and SST-INs expressing iGluSNFR-A184V (green). **B)** Averaged line scan (10 images) of iGluSNFR-A184V fluorescence (green) at contact points between L2/3 PN boutons and PV/SST-IN dendrites after AP (7 APs @ 10 Hz) initiation. Time series traces (10) show mean fluorescence over 1 um (white line), with the black trace as the average (ΔF/F). **C)** Histogram of ΔF/F amplitude distributions for PV (red) and SST-INs (orange) iGluSNFR events. **D)** Summary of failure rates and coefficient of variation for L2/3 PN synaptic contacts in SST-(n=8) and PV-INs (n=9). Data include release probability (Pves) and maximum vesicles released per AP (N max Ves), (*p<0.01, Mann-Whitney test). **E)** Two-photon image of an SST-IN loaded with Alexa 594, with glutamate uncaging at three dendritic locations. Somatically recorded EPSCs from photolysis at two pulse durations. **F)** Similar to **E** for PV-INs. **G)** Summary of paired-pulse ratios of photolysis-evoked EPSCs for PV and SST-INs. **H)** Paired-pulse ratios binned by distance from soma for PV and SST-INs, (**p<0.001, p<0.00001, Kruskal Wallis test with Dunn’s correction). **I, J)** Similar analysis in the presence of NMDA receptor antagonists, alone (I) or with NMDA and VGCC blockers (J), (**p<0.001, p<0.00001, Kruskal Wallis test with Dunn’s correction). **K, L)** Similar analysis for PV-INs with TEA (5 mM), (**p<0.001, p<0.00001, Kruskal Wallis test with Dunn’s correction). Data are mean ± SEM.

We separated successful and failed release events by comparing the amplitude of evoked changes in iGluSnFR fluorescence after each individual-evoked action potential with baseline values (see **Methods** and **Fig. S5**). Overall, as anticipated for low release probability synapses, failures were significantly more pronounced in inputs targeting SST-INs in comparison to those targeting PV-INs (**Fig. 2G,H,I**). These observations were consistent in experiments conducted with both 2 mM (**Fig. 2**) and 1.2 mM extracellular calcium concentrations (**Fig. S5**). In line with the higher number of failures, the coefficient of variation (CV) was greater in inputs onto SST-INs when compared to PV-INs (**Fig. 2C,D and Fig. S5**). The number of release sites (N) defines the maximum number of synaptic vesicles that could be released per action potential at individual synaptic locations (*35*). To estimate N, we performed optical fluctuation analysis (*36*, *37*), assuming a binomial distribution of release events. We found that N was quite variable between individual points of contact, like previously reported in hippocampal INs (*38*), with a mean value per action potential close to 2 for both SST and PV-INs (**Fig. 2D** and **Fig. S5**). Overall, our results reveal that multivesicular release can occur at individual glutamatergic inputs in both cortical SST- and PV-INs.

To test how multivesicular release is integrated in dendrites of PV and SST-INs, we probed the integration of single and double-quanta at single dendritic locations along the dendrites of PV- and SST-INs, using quantal-size adjusted pEPSCs (**Fig. 2E,F**). Dendrites of SST-INs integrated double-quanta linearly all along the dendritic tree (**Fig. 2G,H)**. This linear integration was also observed in current clamp configuration (**Fig. S6**). Conversely, PV-IN dendrites exhibited a progressive somatodendritic attenuation of double-quanta pEPSCs as expected for a passive cable (*13*) (**Fig. 2F-H)**. This decrease in short-term plasticity (STP) was observed also while mimicking depressing inputs that represent the natural plasticity of glutamatergic inputs contacting PV-INs (**Fig. S6**). We subsequently tested the mechanism involved in the relatively linear double-quanta integration along SST-INs dendrites. Blocking NMDARs significantly reduced the ability of SST-IN dendrites to linearly detect two release events at distal dendritic locations (**Fig. 2I,J**). This effect was not further increased by blocking other voltage-gated conductances (**Fig. 2I,J**). In hippocampal PV-INs, activation of potassium channels was proposed to participate in sublinear integration of synaptic inputs (*5*, *39*). However, blocking potassium channels with TEA did not reduce the sublinear integration of double quanta along dendrites in cortical PV-INs (**Fig. 2K,L**).

Our findings indicate that during synaptic transmission, the active dendritic properties of SST-INs contribute to the preservation of presynaptic STP. Conversely, in PV-INs, passive dendrites are expected to dampen STP, particularly for synaptic contacts located within more distal dendritic regions. Consistent with this hypothesis, unitary synaptic responses from paired recordings between L2/3 PNs and PV-INs exhibited a significant inverse relationship between the rise time of EPSCs (a proxy for input location) and short-term plasticity in PV-INs. Such correlation was absent in the unitary EPSCs between PNs and SST-INs (**Fig. S7**). Overall, these experiments suggest that interneuron-specific dendritic operations reduce short-term synaptic plasticity in PV-INs and maintain short-term facilitation in SST-INs. These properties may therefore modify the dynamic activity of PV and SST interneurons during network activity.

## Distinct spatial synaptic distribution along the dendrites of PV- and SST-INs

Our results so far support different dendritic integration properties in neocortical L2/3 PV- and SST-INs. How these operations contribute to integration of synaptic inputs is determined by the timing and location of active synapses. These patterns are partly controlled by the way synaptic inputs distribute along the dendritic tree of neurons. Previous work has shown that synapse density gradients can significantly alter dendritic responses (*40*, *41*) but whether synapse distributions along the dendritic tree are similar between PV- and SST-INs remains unknown. We thus investigated the synaptic distributions along the dendritic tree of PV and SST-INs in L2/3 of S1 (**Fig. 3A**). To estimate the density of glutamatergic synapses we used transgenic mice in which PSD95 (a protein involved in the structure of glutamatergic synapses) was conditionally tagged with a fluorescent marker, mVENUS (*42*, *43*). PSD95 puncta along different dendritic locations in both PV- and SST-INs were imaged using immunohistochemistry and Confocal/STED microscopy. To probe if synapse density was uniform or variable across the dendritic tree, we compared PSD95 puncta density at proximal (< 40 μm from soma) and distal (∼100 μm from the soma) dendritic regions. We found that the densities of glutamatergic synapses greatly varied between proximal and distal locations in PV-INs. The proximal locations exhibited a high density of synapses while the distal locations had a much lower density along individual dendritic branches (**Fig. 3B,C**). On the other hand, the density of synapses did not seem to vary along the dendritic tree in SST-INs (**Fig. 3D-F**).

**Figure 3:**
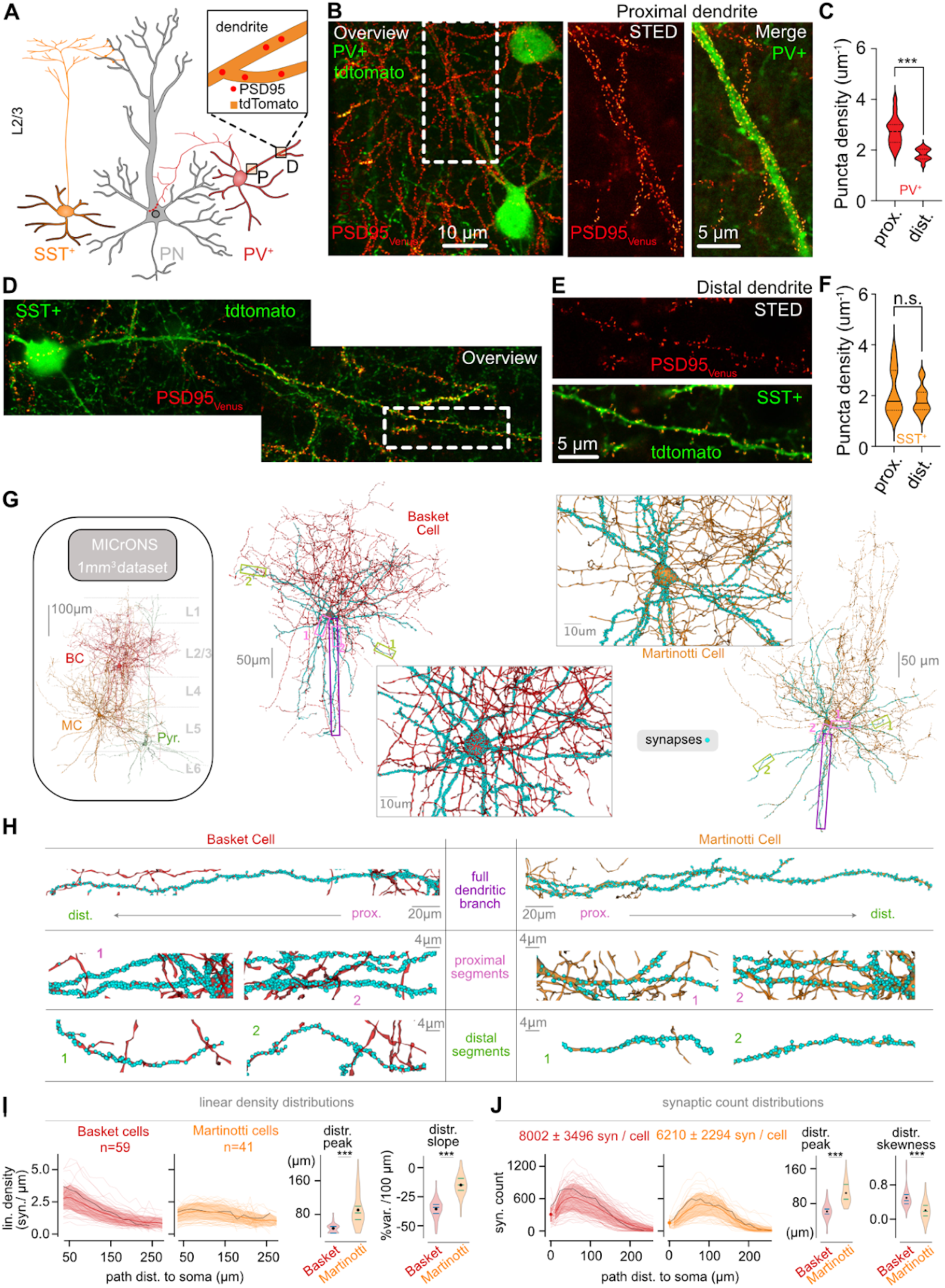
Distinct synaptic distributions in the dendritic trees of Basket and Martinotti INs. **A)** Illustration of experimental approach used to quantify glutamate synapse distribution along dendrites of SST and PV-INs. PSD-95 proteins were selectively labelled in PV- and SST-INs and quantified at two different distances along individual dendritic branches**. B)** (left) Overview image of PV-INs with proximal dendrites labelled with tdTomato (green) and PSD95venus (red). (*Middle*) STED image of dendritic location defined on the left. (*right*) Merged between tdTomato obtained in confocal mode (green) and PSD95venus (red) labelling obtained in STED. **D, E**) Same as **B** but for SST-INs. Note the absence of PSD-95 puncta close to soma but clear labelling in distal dendrites. **C, F**) Summary plot of density of PSD-95 puncta in proximal (less than 40 μm) and distal (approximately 100 μm) dendritic location in PV- and SST-IN dendrites (PV_proximal_, n=33; PV_distal_, n=12, p<0.0001, Mann-Witney test; SST_proximal_, n=21; SST_distal_, n=14, p>0.01, Mann-Witney test). **G**) Example of basket and Martinotti INs reconstructions from the millimeter-scale volumetric electron microscopy MICrONS dataset (*48*). Blue dots indicate the location of identified synapses along the dendrites of a basket and a Martinotti Cell. **H)** Illustration of synapse distributions in individual dendritic branches from reconstructed Basket (l*eft*) and SST-INs (*right*) displayed in **G**. Images display synapses along full individual dendritic branches as well at proximal and distal dendritic segments. **I)** Linear synapse density distributions (*left*, see Methods) for all basket (n=59) and Martinotti (n=41) cells as a function of the distance from soma. (*right*) Violin plots of distribution peaks (Basket vs Martinotti, p=1e-11, Mann-Whitney test) and distribution slopes as a variation per 100μm (Basket vs Martinotti, p=6e-14, Mann-Whitney test) for the plots on the left. **J)** Synaptic count (*left*, see Methods) for all basket (n=59) and Martinotti (n=41) cells as a function of the distance from soma. (*right*) Violin plots of distribution peaks (basket vs Martinotti, p=2e-13, Mann-Whitney test) and distribution skewness (basket vs Martinotti, p=1e-8, Mann-Whitney test) for the plots on the left.

We further investigated such differences in synapse distribution using a publicly available electron microscopy dataset corresponding to the full reconstruction of a 1 mm^3^ sample of the mouse visual cortex (*44*). We combined their databases of (1) proof-read interneuron identifications (*45*), (2) 3D reconstructions of single cell morphologies (*44*) and (3) cell-to-cell synapse identifications (*46*) to estimate synaptic distributions along the dendritic tree of INs with great spatial resolution (**Fig. 3G,H**, see Methods). In this dataset, inhibitory neurons (INs) are morphologically classified, allowing us to focus on basket cells and Martinotti cells as representative proxies for parvalbumin-expressing (PV) and somatostatin-expressing (SST) INs, respectively (*2*, *47*). Consistent with our PSD95 quantifications, the density of synaptic contacts in basket cells gradually decreases with increasing distance from the soma. The synaptic distribution peaks in the very proximal regions, exhibiting a strongly negative slope and highly skewed distributions, **Fig. 3H-J**). On the other hand, the distribution of synapses along the dendritic tree of Martinotti cells was relatively constant along dendritic branches, with distal regions displaying similar densities as compared to proximal regions (medial peak location, negligible slope and weakly-skewed distributions, (**Fig. 3H-J**). Therefore, the major functional dendritic integration properties in PV- and SST-INs described in Figures 1-2 are associated with remarkable distinct synapse distributions, suggesting two markedly different morpho-functional organization principles in these two prominent IN subtypes.

## Two different strategies to enhance EPSP-spike coupling in thin dendritic structures

We next used computational modelling to examine how the two specific dendritic strategies (proximally-biased synapse distributions + low synaptic NMDAR content vs. uniform synapse distribution + higher density of synaptic NMDAR) influence integration of synaptic inputs. We hypothesized that integration properties and synaptic distributions can generate unique functional features of PV- and SST-INs.

For this purpose, we built detailed biophysical models of PV- and SST-INs based on the morphological reconstructions of basket and Martinotti cells respectively, and their own synaptic locations from EM data shown in **Fig. 3G,H**. Synaptic and biophysical properties were constrained by experimental measures (**Table S1-3**, **Methods**). For each dendritic integration strategy, we looked for an alternative “null” hypothesis. In the case of SST INs, this was naturally provided by the removal of the NMDAR component in synaptic transmission. For the PV-IN integration strategy, we generated a uniform surrogate of its synaptic distribution by redistributing synapses on each branch according to a constant linear density. This operation simulated a uniform distribution of glutamatergic synaptic inputs onto dendrites of PV INs. We found that only PV-INs exhibited significant deviations between the real distribution and the uniform surrogate (**Fig. 4A, Fig. S8**).

**Figure 4:**
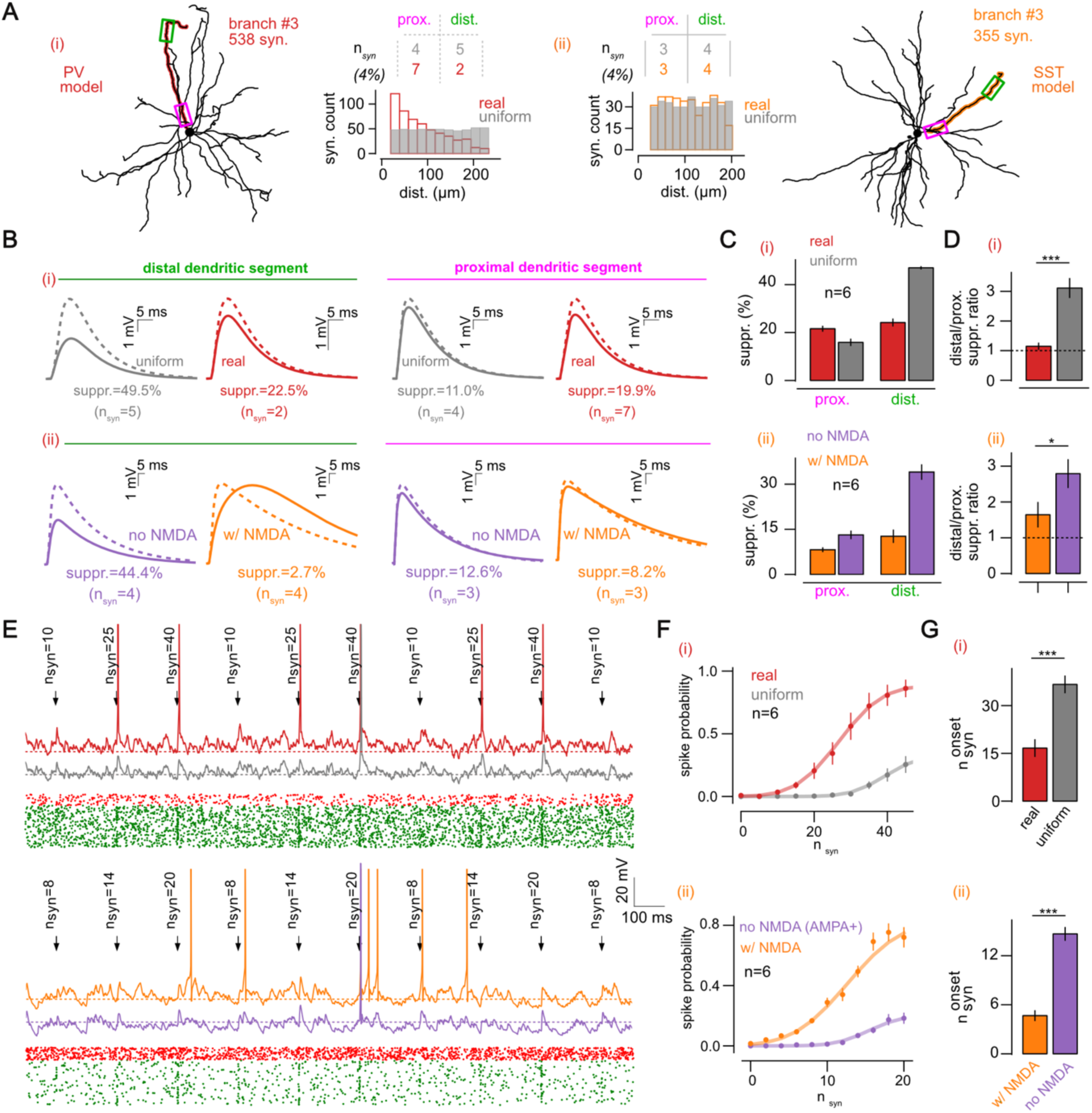
The PV- and SST-IN dendritic programs improve distal signal transmission and optimize synaptic efficacy through different mechanisms. **A)** Morphology of the EM reconstructed basket cell (i) and Martinotti cell (ii) used as PV- and SST-IN models, respectively. In the insets, we show the real distributions (colored bars) of synapses on a single representative branch together with the uniform surrogate distribution (plain gray bars). The inset table shows the sparse subset of synapses in the distal (green) and proximal (pink) segments. **B**) Example numerical simulations of *V_m_* response following the quasi-synchronous stimulation of the synapses in the distal (green, left) and proximal (pink, right) segments in the PV-IN (i) and SST-IN (ii) models. We show the observed response (plain line) and the expected response (dashed line) from the individual event responses. We compare each model to its control situation (i, grey, uniform distribution), (ii, purple, no-NMDA). **C)** Suppression of EPSP amplitude between observed response and linear predictions (see annotations in **B**) in the PV-IN (i, red) and SST-IN (ii, orange) models with their own control in the distal (green) and proximal (pink) segments. **D)** Ratio of suppression between the distal and proximal segments in the PV-IN model (i, red, with its “uniform distribution” control in grey, n=6 branches, p=3e-2, Wilcoxon test) and SST-IN model (ii, orange, with its “no NMDA” control in purple, n=6 branches, p=3e-2, Wilcoxon test). **E)** V_m_ dynamics at the soma following background and stimulus-evoked activity in the dendritic branch shown in **A**. Synaptic stimulation consists in the quasi-synchronous activation of increasing number of synapses *n_syn_* (see annotations). We show the PV-IN model (red, with its “uniform distribution” control in grey) and SST-IN model (orange, with its “no NMDA” control in purple). **F)** Summary plot of the spike probability (in a 50ms post-stim. window) as a function of the number of synapses *n_syn_* in the stimulus. **G)** Onset response level 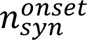 in the PV-IN model (red, with its “uniform distribution” control in grey, p=3e-2, n=6 branches, Wilcoxon test) and SST-IN model (orange, with its “no NMDA (AMPA+)” control in purple, p=3e-2, n=6 branches, Wilcoxon test). Onset level 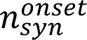 is defined as the input level where the “Erf” fit goes above spike-proba=0.05. Results are shown as mean ± s.e.m over n=6 branches in panels **C**,**D**,**F**,**G**.

In our glutamate uncaging experiments, we observed sublinear integration in PV-INs, while sublinear integration in SST-INs was revealed only upon blocking NMDARs. Based on these findings, we hypothesized that proximally-biased synaptic distribution or the presence of NMDARs might compensate for sublinear integration, as predicted by cable theory (*33*, *48*). This would increase synaptic efficacy and ultimately enhance EPSP-to-spike coupling. To test this, we generated input patterns by randomly picking a sparse subset of synapses (4%, see **Fig. S9**) located on a distal (160-200 μm) and proximal (20-60 μm) segment of a single dendritic branch (**Fig. 4A**) and simulated the arrival of spatially clustered synaptic inputs that are known to engage sublinear integration. This resulted in a small number of synapses (from 2 to 8) that we activated independently and quasi-synchronously (0.5ms delay) to compute the “linear” and “observed” responses in each of the segments (**Fig. 4B**).

For PV-INs, the surrogate, uniform synapse distribution exhibited the distance-dependent sublinear behavior expected for dendrites acting mostly as passive cables (**Fig. 4B**, grey). In the “real” scenario with proximally biased synapse distribution, however, the decreased density of distal synapses resulted in fewer synapses within the 4% cluster (see n_syn_=2 vs n_syn_=5 in the “uniform” surrogate, **Fig. 4A**), preventing synaptic saturation. This is evidenced by a significant reduction in sublinear summation in distal segments (**Fig. 4B-C*i***). On the other hand, the increased number of synapses in proximal segments (in the “real” distribution case compared to the “uniform” surrogate) did not strongly increase sublinearity (**Fig. 4C*i***). Therefore, assuming PV- INs receive signals that are randomly distributed over the dendritic branch, proximally-biased synapse distributions can reduce sublinear dendritic integration in distal dendritic segments and limit signal loss by reducing the size of active synaptic clusters (**Fig. 4D*i***).

As expected, in the case of SST-INs, the “no-NMDA” surrogate also exhibited a distance-dependent sublinear behavior with strong distal suppression and near-linear integration in proximal dendritic segments (**Fig. 4B*ii***, purple). Activation of NMDARs completely abolished this effect during multi-input integration (**Fig. 4C*ii***) due to the involvement of the slow NMDAR excitatory conductance (**Fig. 4B*ii***, orange). Furthermore, in agreement with our glutamate uncaging results, increasing the number of active synapses during multi-input integration (especially in distal dendritic segments) resulted in supralinear integration (**Fig. S9**). We conclude that the presence of a strong NMDAR component observed in the dendritic program of SST-INs not only limits sublinear integration, but it also amplifies coincident synaptic activity along dendrites, especially in distal dendritic locations, like PNs (**Fig. 4D*ii***).

Finally, we analyzed the impact of the differential anatomical and functional synaptic strategies exhibited by PV and SST cells on firing dynamics under a more realistic setting. Specifically, we tested the ability of synaptic stimulation to elicit action potentials on a *per branch* basis. Neuronal activity was simulated in response to increasing synapse number *n_syn_* (randomly picked on the branch), on top of a stationary excitatory and inhibitory background to mimic *in vivo*-like activity (**Fig. 4E**, see Methods). PV-INs with real synaptic distributions were substantially more effective in driving spiking activity than with uniform distributions (**Fig. 4F,Gi**). Likewise, SST-INs exhibited a substantial decrease in the spiking probably in response to synapses devoid of NMDARs (**Fig. 4F,Gi**, despite a boost of the AMPAR conductance weight to match EPSP size at -60 mV, see **Methods**).

We conclude that the differential morpho-functional dendritic integration strategies exhibited by L2/3 PV- and SST-INs represent two distinct strategies to increase synaptic efficacy. Synapses on PV-INs are positioned strategically to reduce the likelihood of sublinear integration, especially in distant dendritic areas. Conversely, SST-INs take advantage of the NMDARs to enhance coincident dendritic activity all along their dendritic arbor.

## Different dendritic integration strategies in PV- and SST -INs result in temporally-distinct output firing dynamics

While both dendritic integration mechanisms can compensate for the synaptic strength shortcomings of sublinear summation, the slow NMDA receptor conductance is likely to produce a very different temporal profile of IN spiking (*49*). Indeed, in our detailed biophysical models, a striking difference between PV- and SST-INs was their specific temporal window for input selectivity and output spike patterns (**Fig. 5A**). The spiking pattern of the PV-IN model accurately transmitted the amplitude and width of step stimulations (**Fig. 5A**, red, note the similarity between input steps and output rate responses) while the SST-IN model would produce a more integrative and long-lasting IN-spiking response due presence of the slow NMDAR-mediated conductance (**Fig 5A**, orange, note the equal rate levels in the second step despite a twice weaker amplitude, note also the delayed responses outside the stimulus window). To generate testable predictions, we simulated a more realistic input pattern by stimulating the IN models with temporally correlated excitatory and inhibitory activity (**Fig. 5B**, the input strength was scaled so that average output firing lies in the 10-15 Hz range, see **Methods**). We then analyzed the temporal dynamics of the two IN models by computing the cross-correlation function of its output spiking rate (after trial averaging) with the input signal (in grey, in **Fig. 5A**). The fast and linear report of response duration of PV-IN resulted in a sharp inhibition closely following the input temporal correlations (**Fig. 5C**, captured by a sharp cross-correlation half-width in **Fig. 5D**). The broader and long-lasting responses of SST INs resulted in a delayed and extended firing dynamics (**Fig. 5C**, captured by a broader cross-correlation half-width in **Fig. 5D**). These initial predictions are consistent with the known coincidence detection mode of PV-INs and the more integrative mode of SST-INs (*16*).

**Figure 5:**
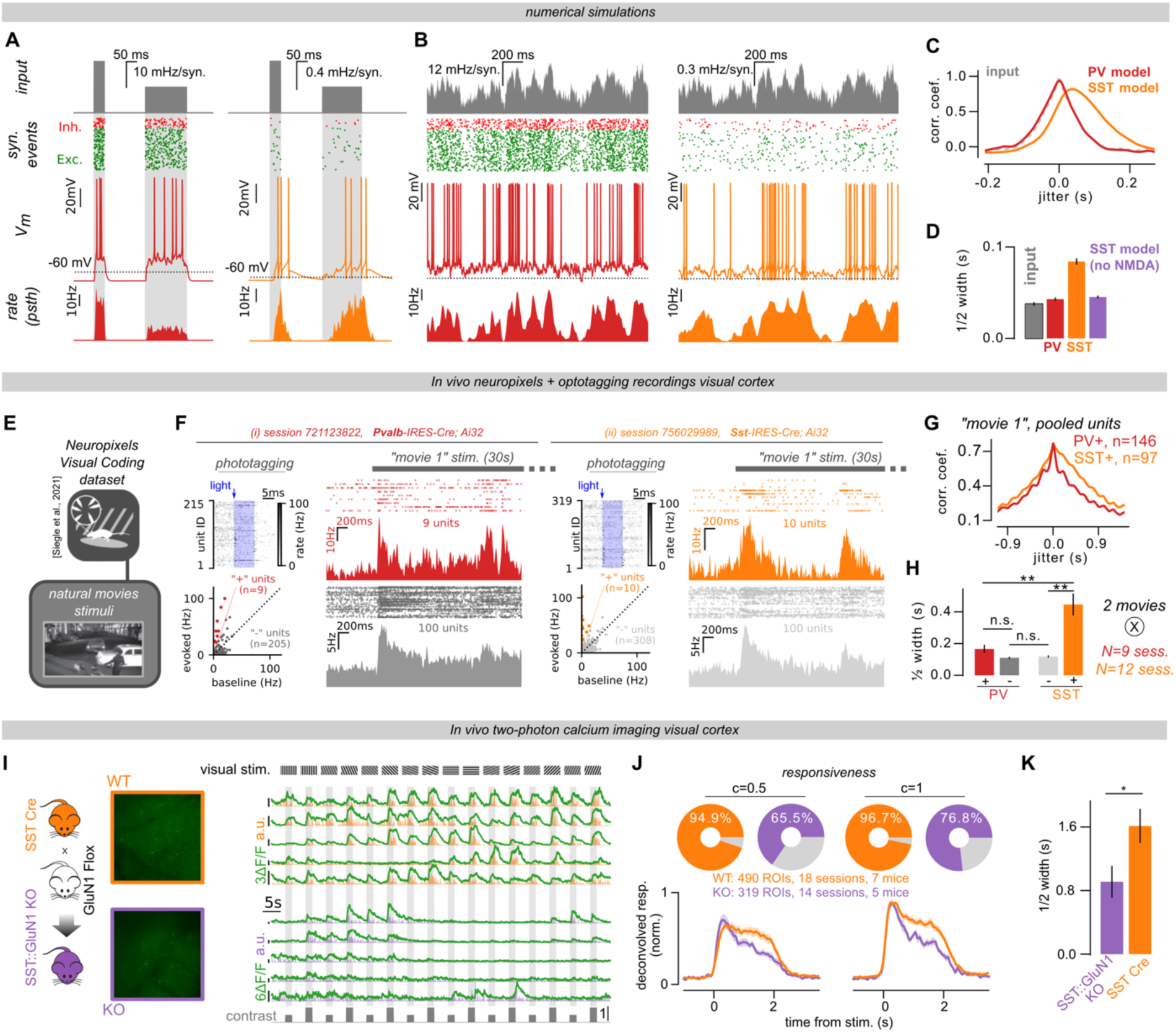
The PV- and SST-IN dendritic programs shape two temporally-distinct inhibitory dynamics in cortical networks. **A)** Response to packets of excitatory and inhibitory activity controlled by current steps in the PV-IN model (red) and in the SST-IN model (orange). We show the time-varying rate (top, grey) controlling the generation of excitatory and inhibitory events (green and red respectively), a single trial V_m_ example (middle) and the time-varying output rate bult from multiple trials (n=24). **B**) Same than **A** but for a time-varying input rate controlled by a temporally-correlated stochastic process (see Methods). **C)** Cross-correlation function between input rate and PV-IN (red) and SST-IN (orange) rate. We also show the autocorrelation function of the input rate (grey). **D)** Positive half-width of the cross-correlation functions (see Methods, shown as mean ± s.e.m. over N=4 input seeds x n= 6 branches; input vs PV, p=6e-3; input vs SST, p=3e-9; input vs SST-noNMDA, p=3e-3; PV vs SST:, p=3e-9; SST vs SST-noNMDA, p=3e-9; Mann-Whitney test). **E)** Schematic of the Neuropixels dataset from (Siegle et al., 2021). **F)** Single session examples in PV-cre and SST-cre (ii) mice. We show the photo-tagging trials (top-left) and the summary analysis to split positive and negative units (see Methods). We show the spiking events of positive (colored) and negative (grey) units around stimulus onset (right). **G)** Cross-correlation function between negative unit rates and PV-positive rate (red) or SST-positive rate (orange) for the natural movie #1 shown in all sessions. **H)** Half-width of the cross-correlation function in the positive and negative units. PV-cre mice: N=9 sessions x 2 movies, PV+ units vs PV-units, p=7e-2, Mann-Whitney test. SST-cre mice: N=12 sessions x 2 movies, SST+ units vs SST-units, p=2e-5, Mann-Whitney test. PV-units vs SST-units: p=5e-1, Mann-Whitney test. PV+ units vs SST+ units: p=3e-3, Mann-Whitney test. Results are shown as mean+/-sem over sessions and movies. **I)** (left) Illustration of the genetic mouse model approach to selectively remove NMDARs from SST-INs. (Middle) Two-photon image of SST-INs in V1 expressing GCaMP6s. (Right) Representative fluorescence traces shown as ΔF/F0 (green) together with their deconvolution traces. **J)** Percentage of SST-INs exhibiting a statistically significant positive response to visual stimuli at both full and half contrast in wild-type subjects and in animals lacking GluN1 subunits selectively in SST-INs. **K)** Deconvolved responses following stimulus presentation (average over all orientations) at half and full contrast. In the inset, we show the value of a Gaussian curve decay in the fitting window highlighted in grey. **L)** Half-width of the evoked response decay (evaluated by the width parameter of a Gaussian fit in the window highlighted in grey). SST:WT vs SST:GluN1-KO, p=4e-2, Mann-Whitney test.

We tested the above predictions in a publicly available dataset of single unit recordings from the mouse visual cortex in response to natural movies presentation, in which PV- and SST-IN firing was identified by photo-tagging (*50*) (**Fig. 5E**, see **Methods**). Specifically, we computed the time-varying firing rate of each IN type by pooling the positive units of a single session (n=24±14 per session, N=9 sessions for PV-INs, n=8±6 per session, N=12 sessions for SST-INs). Because both PV- and SST-INs are primarily driven by local cortical excitation (*51*), we estimated the input signal by computing the time-varying rate of 100 randomly picked, non-photo-tagged (negative) units in the recordings, assumed to originate from putative PNs (**Fig. 5F**). We next computed the cross-correlation between input (from negative units) and IN rates (from positive units) (**Fig 5G)**. In accordance with our model predictions, we observed a wider temporal window in SST INs in the cross-correlation function, as compared to PV INs (**Fig. 5G**). This observation was highly consistent across sessions and movies, as measured by the half-width of the cross-correlation function (**Fig. 5H**). We conclude that the population dynamics of PV- and SST-INs have temporal features compatible with our observed distinct dendritic integration strategies.

Importantly, PV- and SST-INs do not vary only in their dendritic integration strategies, and therefore other mechanisms could underlie these differences. For example, synaptic facilitation at the excitatory to SST-INs synapses could also contribute to such extended temporal features. To specifically address this concern, we tested a more specific prediction of the model: the wider and delayed responses of SST-INs should be significantly reduced upon removal of NMDARs from this IN subclass (**Fig. 5C**). To this purpose, we conditionally knocked out NMDARs by crossing GluN1^fl/fl^ with SST-Cre mice (SST:GluN1-KO mice). Importantly, despite lacking NMDARs, glutamatergic synaptic transmission and short-term plasticity at L2/3 PNs SST-INs synapses were not altered in SST:GluN1-KO mice (**Fig. S10**). We then expressed the genetically encoded Ca^2+^ sensor GCaMP6s in SST INs and recorded their activity in V1 of both control mice and mice lacking NMDARs specifically in SST-INs (**Fig 5I** and **Methods**). Fluorescence signals were deconvolved to remove the calcium-sensor dynamics (see **Methods**). Awake mice were presented full-field grating stimuli at two contrasts (0.5 and 1) and 8 orientations. We analyzed the properties of the stimulus-evoked activity (**Fig. 5I**). In control conditions, SST-INs responded robustly to visual stimuli (**Fig. 5J**). We observed a significant decrease of SST-INs responsiveness at half and full contrast in SST:GluN1-KO mice, thus supporting the role of the NMDARs in shaping SST-INs activity (**Fig. 5J**). In agreement with the results of **Fig. 5A-H**, visually evoked responses of SST INs exhibited much sharper temporal dynamics in SST:GluN1-KO mice as compared to their WT littermates (**Fig. 5K**). Our findings indicate that the SST-IN-specific dendritic strategy to integrate glutamatergic synapses using NMDARs plays an important role in shaping the temporal features of SST-mediated firing behavior in response to visual stimuli.

## Discussion

In this study, we provide an in-depth analysis of the dendritic integration properties of two major inhibitory neuron classes, PV- and SST-INs, which are characterized by distinct connectivity properties and functional roles within cortical networks (*2–4*). Despite the extensive body of evidence examining dendritic integration in cortical PNs, little is known about how synaptic activity is processed by dendrites of cortical INs. We found that PV- and SST-INs use two different strategies to integrate inputs along their dendritic trees. Interestingly, our model simulations predict that variations in the density of synaptic NMDARs is a major contributor for such differential dendritic integration properties. Previous studies reported that the density of NMDARs is quite variable across IN classes (*49*, *52*), but the functional relevance of such different expression levels of NMDARs was unclear. We propose that differences in the content of synaptic NMDARs across populations of interneurons contribute to generate functional diversity in their dendritic operations and ultimately define their role during *in vivo* network activity. Such a hypothesis is also supported by recent findings in PNs where variability of NMDA-to-AMPA ratio was reported to result in the presence of different modes of dendritic integration in L5b PNs of the retrosplenial cortex to for distinct long-range inputs (*53*).

The present results suggest that, in addition to the known morphological, transcriptomic, and electrophysiological differences, dendritic integration properties vary across the distinct populations of cortical interneurons. Numerical simulations highlight that such a variability in integration properties plays a crucial role in shaping the distinctive functional characteristics of specific interneuron classes, particularly PV- and SST-INs. In PV-INs, dendrites behave mostly as passive cables, due to the combination of reduced NMDAR levels, and low input resistance. This leads to a synaptic integration mechanism that enables rapid membrane voltage deflections in response to increased excitatory synaptic activity. Such a dendritic performance allows precise temporal response of PV-INs to variations in pyramidal cell activity. These properties seem to be particularly well tailored for PV-INs to fulfill their important role as sharp coincidence detectors, to sharpen sensory inputs through powerful and precise feedforward inhibitory networks, and drive fast cortical brain oscillations (*18*). Conversely, in SST-INs, the presence of slow NMDARs along dendrites extends the window of temporal integration, the efficacy of synaptic activity in driving SST-INs firing. This extended temporal window appears important for providing feedback inhibition to cortical microcircuits using glutamatergic synapses that are known to be particularly inefficient because of low release probability (*19*, *20*).

Surprisingly, we found that the different dendritic integration modes of PV and SST INs were associated with different distributions of glutamatergic inputs along their dendritic trees. Ultimately, the engagement of dendritic operations relies on the spatial and temporal patterns of activation of many synaptic inputs. Currently, information about the patterns of synaptic activity reaching dendrites of PV- and SST-INs *in vivo* is lacking. Yet, our examination of the distribution of synapses reveals a non-uniform pattern along the dendritic branches of PV-INs. This raises the hypothesis that, during cortical processing, the number of local simultaneously active synapses (active clusters) along PV-IN dendrites may be variable, which could have an impact on non-linear dendritic engagement. In the cerebellum, gap junctions have been reported to compensate for sublinear integration (*14*). Here, we propose that gradients of synapse distributions in PV INs may have a similar effect by reducing the number of coactive synapses in distal dendritic regions, where input resistance is particularly elevated, and dendritic saturation is easily achieved. Interestingly, in dendrites of SST-INs where synapses contain higher density of NMDARs, this effect was much less pronounced. Altogether, our data suggest a general scenario in which INs use a variety of dendritic integration strategies to enhance the efficiency of synaptic transmission into post-synaptic spikes (*54*), but tailor the timing of that activity to the specific functional attributes of each IN population.

The asymmetric distribution of synapses that we report here does not preclude the occurrence of sublinear integration in the dendrites of PV-INs. Ultimately, the clustered activation of inputs depends on the spatial and temporal arrival of synaptic activity in dendrites *in vivo*, determined not only by the hardwiring inputs but also by their timing of activation. Non-linear dendritic operations, particularly in PV-INs in the hippocampus, have been proposed to be important for enhancing memory encoding and to detect network ripple activity in PV-INs (*9–12*, *55*). However, such diverse functions of PV-INs require precise hardwiring and spatio-temporal organization of inputs into PV-INs to contribute to the reported physiological phenomena. Here, we propose that, in a more general framework without assuming a specific connectivity motif, PV-INs bias their input distribution to proximal regions of the soma to maximize EPSP-to-spike coupling and reduce saturation in the distal parts of their dendrites. Nevertheless, future studies are crucial to dissect the physiological patterns of synaptic activity that occur in PV-as well as in SST-INs. Additionally, in our study, we primarily focused on studying the integration of voltage signals. It is known that calcium signaling might not necessarily follow the same integration path as the synaptically-evolved EPSP (*6*).

Overall, we describe two simple yet clever strategies employed by PV- and SST-INs to prevent synaptic saturation at their distal dendrites: removal of NMDARs and concentration of glutamatergic synapses towards the soma in PV-INs, and enrichment of NMDARs all along the dendrites in SST-INs. These differential strategies result in profoundly different transformations of excitatory recruitment signals into firing activity by each IN subclass, ultimately shaping their distinct roles within cortical networks.

## Acknowledgements

This study was supported by the Centre National de la Recherche Scientifique “Investissements d’avenir” ANR-10-IAIHU-06, Agence Nationale de la Recherche (to N.R.; ANR-16-CE16-0007-02; ANR-18-CE16-0011-01; ANR-20-CE16-0011-01; ANR-22-CE37-0008-01 to A.B.; ANR-DecoSensoMol1 to N.R. and A.B.; ANR-17-CE16-0026-01 to A.B. and D.D.), ANR-23-CE16-0003 to N.R. and D. D. and ANR-16-CE19-0003-01 to G.M.), the Émergence programme par l’Alliance Sorbonne Université (J.L.); the European Research Council (ERC-STG-678250 to N.R.), the European Union’s Horizon 2020 research and innovation programme (Marie Skłodowska-Curie Grant 892175 “*InProsMod”* to Y.Z.); by the Paris Brain Institute, and the Fondation pour la Recherche Médicale (FRM; fellowship ARF201909009117 to Y.Z.); Equipes FRM– DEQ20150331684; Equipes FRM – EQU201903007860 to A.B.) by NIH (1R01MH124867-02) to P.P. by the NeuroCure Cluster of Excellence Berlin, by the Deutsche Forschungsgemeinschaft [DFG], SFB-1315 ‘Brenda Milner Award’ to AT; by Région Ile-de-France (DIM-ELICIT grant to G.M. and N.R.) We thank the Paris Brain Institute core facilities, namely iVector and ICMice PHENOPARC. We are very grateful to Casey Schneider-Mizell, Forest Collman and Emily Joyce for sharing material and guiding us in the analysis of the MICrONs dataset and to Kathleen Cho and Hillel Adesnik for comments on the manuscript.

## Author contributions

Conceptualization: AM,YZ, AB, DD, JL and NR

Methodology: AM,YZ, DhD,EB,AT,GM,LC,PP,AB,DD,JL and NR

Investigation: AM,YZ, DhD,EB,AT,GM, JL and NR

Visualization: AM,YZ, DhD,EB,AT,GM,LC, PP, AB,DD, JL and NR

Funding acquisition: YZ, DD, AB,JL and NR

Project administration: NR Supervision:AB,DD,JL and NR,

Writing – original draft: AM,YZ,AT,PP,AB,DD,JL and NR

Writing – review & editing: AM,YZ,AT,LC,PP,AB,DD,JL and NR

## Data and materials availability

The code for the data analysis (EM dataset, Neuropixels dataset, Imaging dataset) and numerical simulations (simplified morphological model, detailed biophysical and morphological models) of this study is publicly available at the following link: https://github.com/yzerlaut/pv-sst-dendrites.

## Supplementary Materials

**Figure S1:**
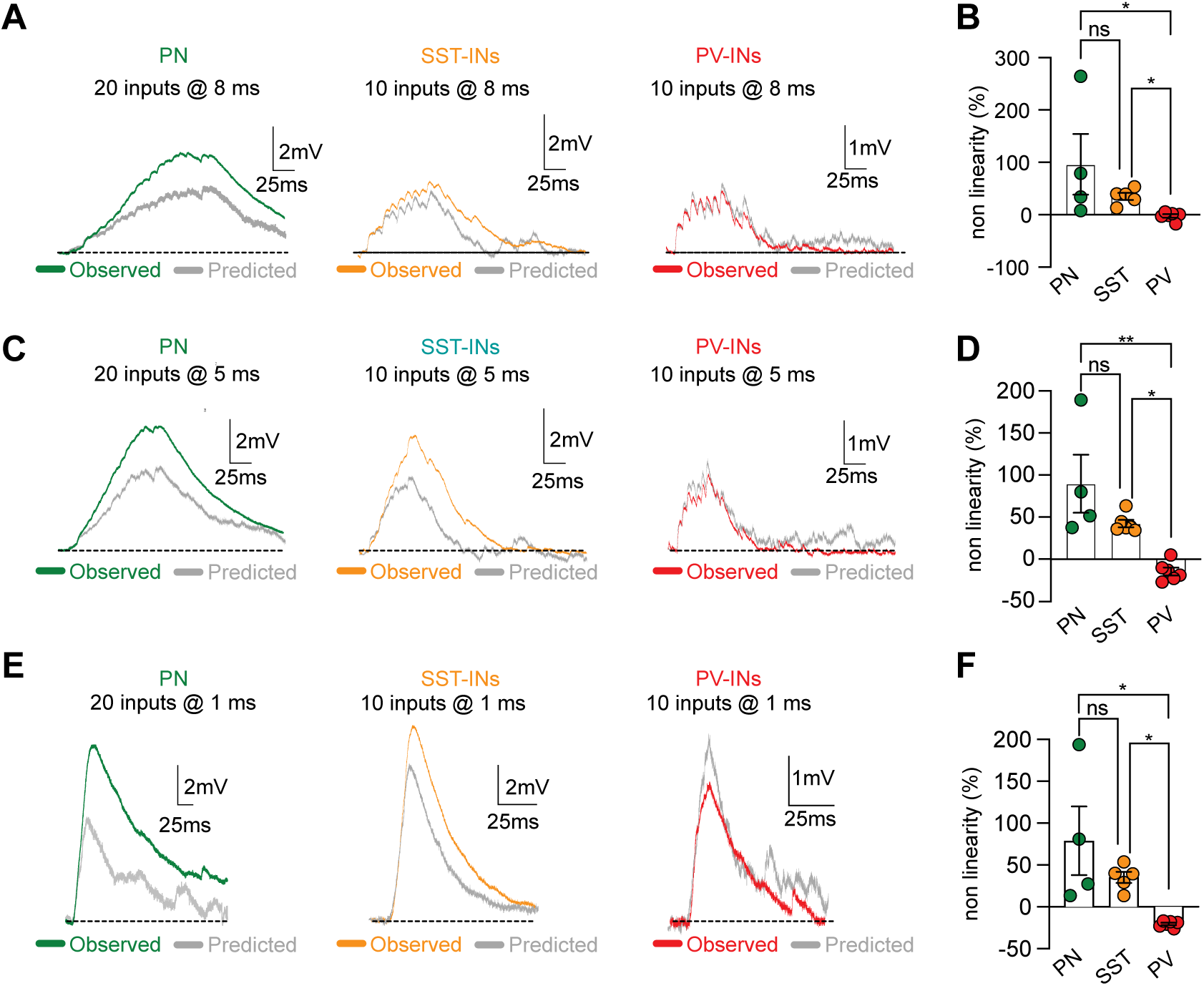
Dependence of synaptic integration in PN, SST and PV-INs on the time interval between uncaging events. **A)** Population average pEPSPs recorded at soma in response to 20 (PN) or 10 uncaging spots (PV and SST-INs) activated with 8 ms inter pulse interval. Recorded average is presented in color while the linear sum is depicted in grey. **B)** Summary plot of non-linearity obtained across the different population of cells using 8ms interpulse interval. Data are presented as average ± SEM with single experiment quantified as dotted color (PN: 96.36±57.88%, n=4; SST: +35.08±6.69%, n=5; PV:-2.32±3.25,n=6; PN vs SST p>0.99; PN vs PV p=0.02; SST vs PV p=0.03; Kruskal Wallis test with Dunn’s correction). **C, D)** Same as A and B, but for 5 ms interpulse interval (PN:89.66±34.39, n=4;SST: 42.36±4.45, n=6; PV:-14.64±4.68, n=6; PN vs SST p=0.83; PN vs PV p=0.003; SST vs PV p= 0.046; Kruskal Wallis test with Dunn’s correction). **E, F)** Same as A and B, but for 1 ms interpulse interval (PN:78.85±41.01, n=4;SST: 30.41±5.83, n=5; PV:-20.49±1.47, n=6; PN vs SST p>0.99; PN vs PV p=0.01; SST vs PV p= 0.03; Kruskal Wallis test with Dunn’s correction).

**Figure S2:**
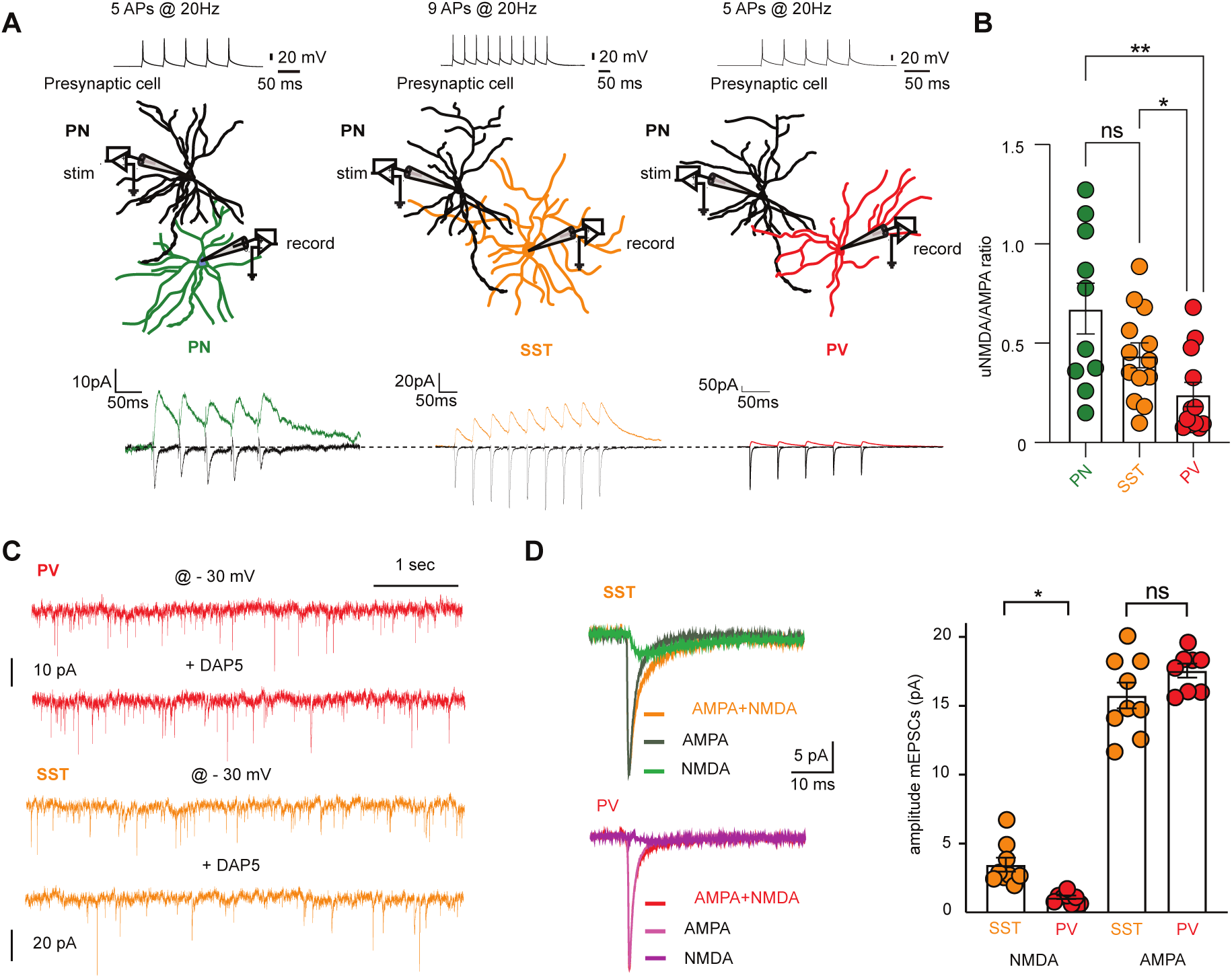
Different synaptic AMPA-to-NMDA receptor ratio in SST and PV-INs. **A)** (*top*) Schematic representation of the experimental approach used to monitor unitary EPSCs between individual L2/3 pyramidal neurons (pyr) and different postsynaptic partners. Unitary AMPA- and NMDA-EPSCs were separately recorded by varying the holding membrane potential of the postsynaptic potential from -70 mV (uAMPA-EPSCs) to + 40 mV (uNMDA-EPSCs). NBQX was added to the recording solution at positive potentials to block AMPA+Kainate receptors. Synaptic currents were recorded in response to a presynaptic train of action potential evoked at 20 Hz through current injection. (*bottom*) Representative AMPA- and NMDA-EPSCs (average of 30 sweeps) obtained for the different postsynaptic partners. **B)** Summary plots for the experiments illustrated in A reported as uNMDA to AMPA ratio (PN n=10; SST n=14; PV n=14). The data are reported as mean ± SEM (PN: 0.67 ± 0.13; SST: 0.44 ± 0.06; PV:0.23 ± 0.05. PN vs SST p=0.94; PN vs PV, p= 0.004; SST vs PV p= 0.04, Kruskal Wallis test with Dunn’s correction **C)** Example traces for miniature EPSCs (mEPSCs) recorded at -30mV in a PV(red) or SST(orange) INs in control conditions and in the presence of NMDA blocker D-AP5. **D)** Average or aligned mEPSC events recorded in both SST and PV cells under control conditions (AMPAR+NMDAR), as well as after bath application of D-AP5 (AMPAR only). The subtraction between these two conditions allowed for isolating the slow component, representing the contribution of NMDAR alone (NMDA). Summary plot of the NMDAR- and AMPAR-mediated components of mEPSCs recorded in SST- and PV-INs determined using the subtraction described on the left. The data are presented as average ± SEM.

**Figure S3:**
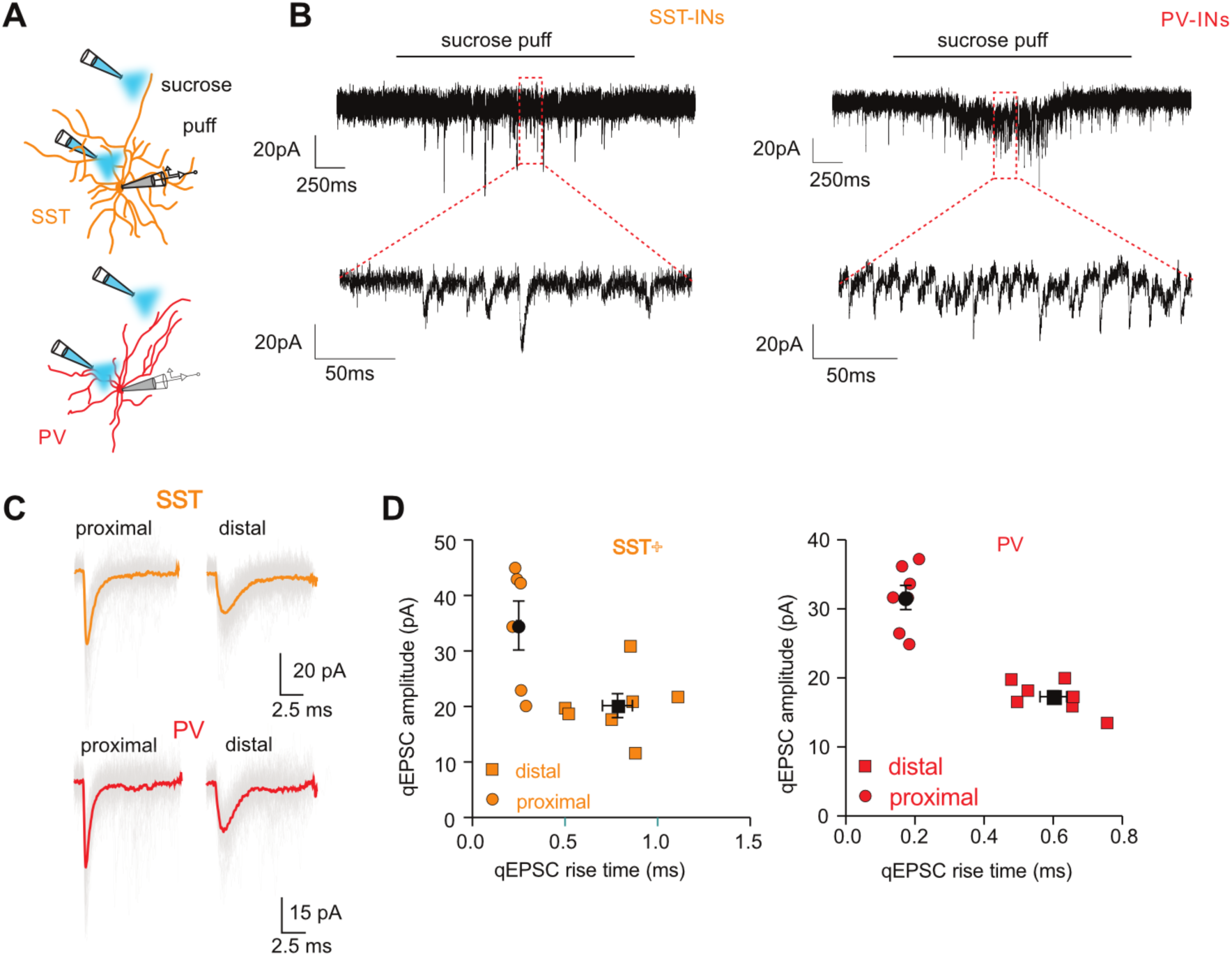
Experimental estimation of single quanta EPSC (qEPSC) amplitude for proximal and distal dendritic locations in SST- and PV-INs. A) Graphical representation of the experiment used to monitor qEPSC at different dendritic locations. Quantal responses were obtained by local perfusion of a hypertonic sucrose solution placed at proximal or distal dendritic locations using two-photon imaging (see Methods). B) *Top:* Example traces of recorded EPSCs during a 3 sec puff application of sucrose. *Bottom:* Magnification of recorded segment showing the sucrose evoked events in SST and PV cells. **C)** Example traces of detected and aligned sucrose-evoked EPSCs. Average mEPSC is shown in color. *Right:* Quantification of the amplitude of average mEPSCs obtained for proximal (square) and distal (circle) dendritic locations in both SST (orange) and PV cells (red) plotted in function of the rise time. The values are reported as color-coded for single experiments with the average and SEM in black (SST proximal: 34.59±4.40 pA; SSTdistal: 20.15±2.18 pA; PVproximal: 31.66±1.74 pA; PVdistal: 17.21±0.83 pA). **D)** Quantification of rise time and amplitude measured after excluding fast events from distal PV recordings using 2SD of the rise time obtained from proximal recordings (see Methods). Plots are presented as mean±SEM with change in single value reported as a horizontal bar (rise time all: 0.45±0.05; selected: 0.59±0.04, p=0.01 Wilcoxon test; Amplitude all:17.69±0.84; amplitude selected: 17.16±0.82, p=0.30, Wilcoxon test.

**Figure S4:**
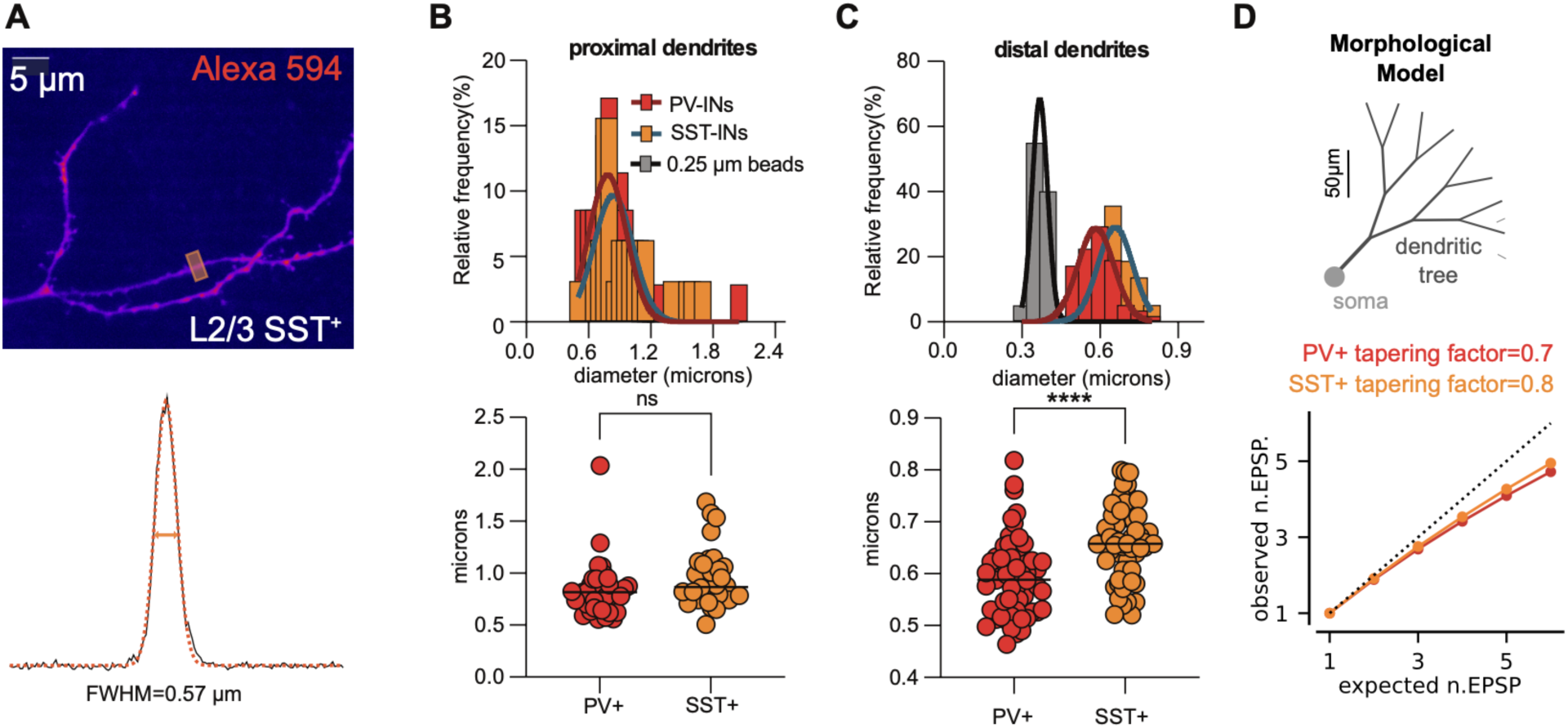
Dendritic diameters can not explain the qualitative difference in dendritic integration properties between PV- and SST-INs. **A)** Example of a 2P image capturing SST-INs distal dendritic branches. The orange square defined the dendritic location used to create the intensity profile shown and quantified below (see methods). **B)** *Top:* Distribution of dendrite diameters for proximal (<40μm) PV-(red) and SST-INs (orange) dendritic segments. *bottom:* Summary plot of average proximal dendrite diameters obtained for PV and SST-INs . The data are presented as individual values and population median (black line). PV median:0.82 mm, n=35; SST median:0.87 mm, n=32; p=0.07 Mann Whitney test. **C)** *Top:* Same as in B but for distal (>100μm) dendritic locations. In gray the distribution for the detected diameter of 0.25 um beads, used to inform about lateral resolution of the 2P system. *Bottom:* Same than in **B** but for distal dendritic locations. The data are presented as single values and the median (black line). PV median: 0.59 mm, n=58; SST median: 0.66 mm, n=59; p<0.0001 Mann Whitney test. **D)** Variations of dendritic diameters in the range of the experimentally-measured differences of **C** predicts minor impact on the dendritic integration mode in the simplified morphological model (see Methods). The root diameter of the morphological model is kept identical between the PV and SST model following the data of **B**. The tapering factor (the factor between the diameter of the children branches and the parent branch) was increased by 12% following the data of **C**. The average response over the 4 locations shown in Figure 1I is shown. The change in non-linearity between the PV and the SST morphologies is 2.3%.

**Figure S5:**
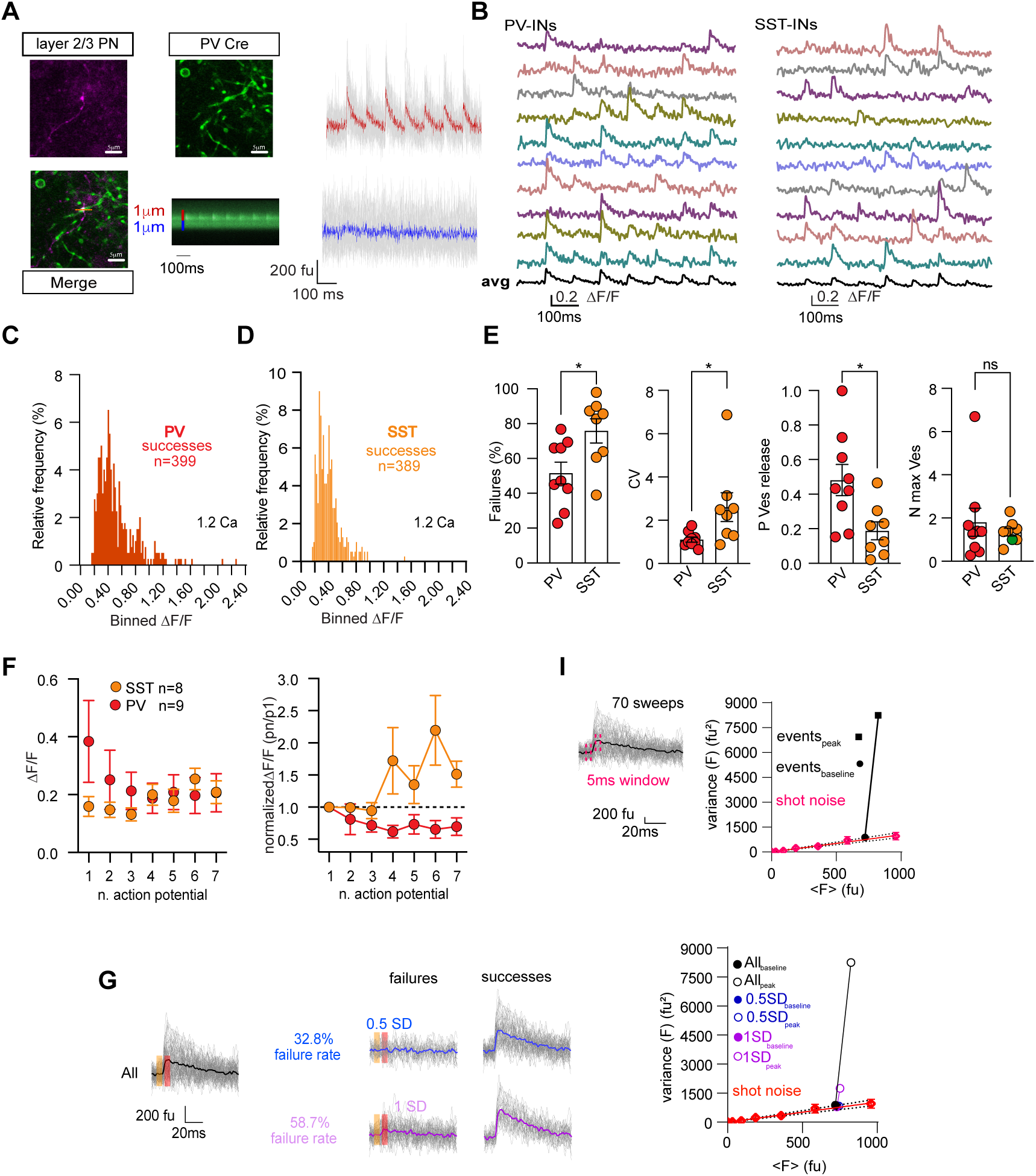
Quantification and analysis of iGluSnFR fluorescence events. **A)** *Upper left:* MIP of a L2/3 PN axon patch loaded with Alexa 594. *Upper middle:* Positive iGluSnFR dendrites present in the same field of view as the image on the left. iGluSnFR was selectively expressed in PV-INs using PV-Cre transgenic mice*. Bottom left:* Point of contact between the Alexa 594 fluorescence (axon) and iGluSnFR signal (putative PV-dendrite) where release of glutamate was monitored using 2P linescan. *Bottom middle:* Average 2P linescan (10 sweeps) obtained from point of contact illustrated in merged image. The increase in iGluSnFR fluorescence observed upon presynaptic firing of the individual L2/3 PYRs is restricted to a narrow (∼ 1 μm) dendritic space. *Right:* Time series traces are mean fluorescence of individual images over 1 μm (red and blue line), and the trace is the average of all 10 traces. **B)** Single trial iGluSnFR fluorescence signal after conversion to DF/F in a PV(red) and a SST(orange) postsynaptic dendrite in the presence of 1.2 mM extracellular Ca^2+^. In black the average of 10 sweeps. **C)** Histogram of DF/F amplitude distributions of all iGluSnFR events detected in PV(orange) recorded at 1.2 mM extracellular Ca^2+^(bin value=0.03 DF/F). **D)** Same as **C** but for SST dendrites. **E)** Summary plots of failure rates and coefficient of variation obtained from iGluSnFR imaging in 1.2 mM extracellular calcium for PV-(*n* = 9) and SST-(*n* = 8) INs. Release probability and number of release vesicles per action potential were calculated using optical fluctuation analysis in the 1.2 mM Ca^2+^ condition assuming a binomial distribution model. The data are presented as mean±SEM (Failures PV: 51.64±6.30%; SST: 75.94±6.94%, p=0.04; CV PV: 1.11±0.11; SST: 2.61±0.66, p=0.01; P Ves release PV: 0.48±0.09; SST: 0.19±0.05, p=0.01; N max Ves PV: 1.80±0.65; SST: 1.35±0.19, p=0.91. Mann-Whitney test. **F)** *Left:* Mean ± SEM of the DF/F values obtained per AP. *Right:* normalized DF/F values from the experiments on the right. **I)** Example of iGluSnFR fluorescence recorded for 70 APs (grey). Average trace is shown in black. The dotted line represents the time interval (5ms) and the location in the events used for measuring baseline and peak fluorescence values. On the right the graphical representation of the mean-variance plot for the baseline (black circle) and the peak fluorescence values (black). The magenta circle represents the mean-variance plot obtained by imaging the patch pipette at different laser powers to estimate variance expected from shot noise. In dotted black the 99% confidence interval for the measurement (see methods). **G)** *Left:* Example of iGluSnFR fluorescence signals recorded for 70 APs (grey) evoked in a single presynaptic L2/3 PYRs. Average response is shown in black. The two colored 5 ms windows represent the location for the baseline and peak fluorescence measurements. *Center:* Example of failure rate detection obtained using two different amplitude thresholds. The success detection was performed using a thresholding method comparing amplitude in peak window + SD of baseline (baseline average±0.5 or 1 SD of the baseline; see methods). Different multiple factors of SD were used and best value was chosen using mean-variance plots like illustrated in right. *Right:* Mean-variance plot for baseline and peak windows on the traces illustrated in the left using baseline± 0.5 or 1 SD. The empty circles represent the peak values and the full circle the baseline. Based on the peak variance detection the value to be used for the failure detection is baseline ± 0.5 SD. In this case the experiment presents a failure rate of 32.8%.

**Figure S6:**
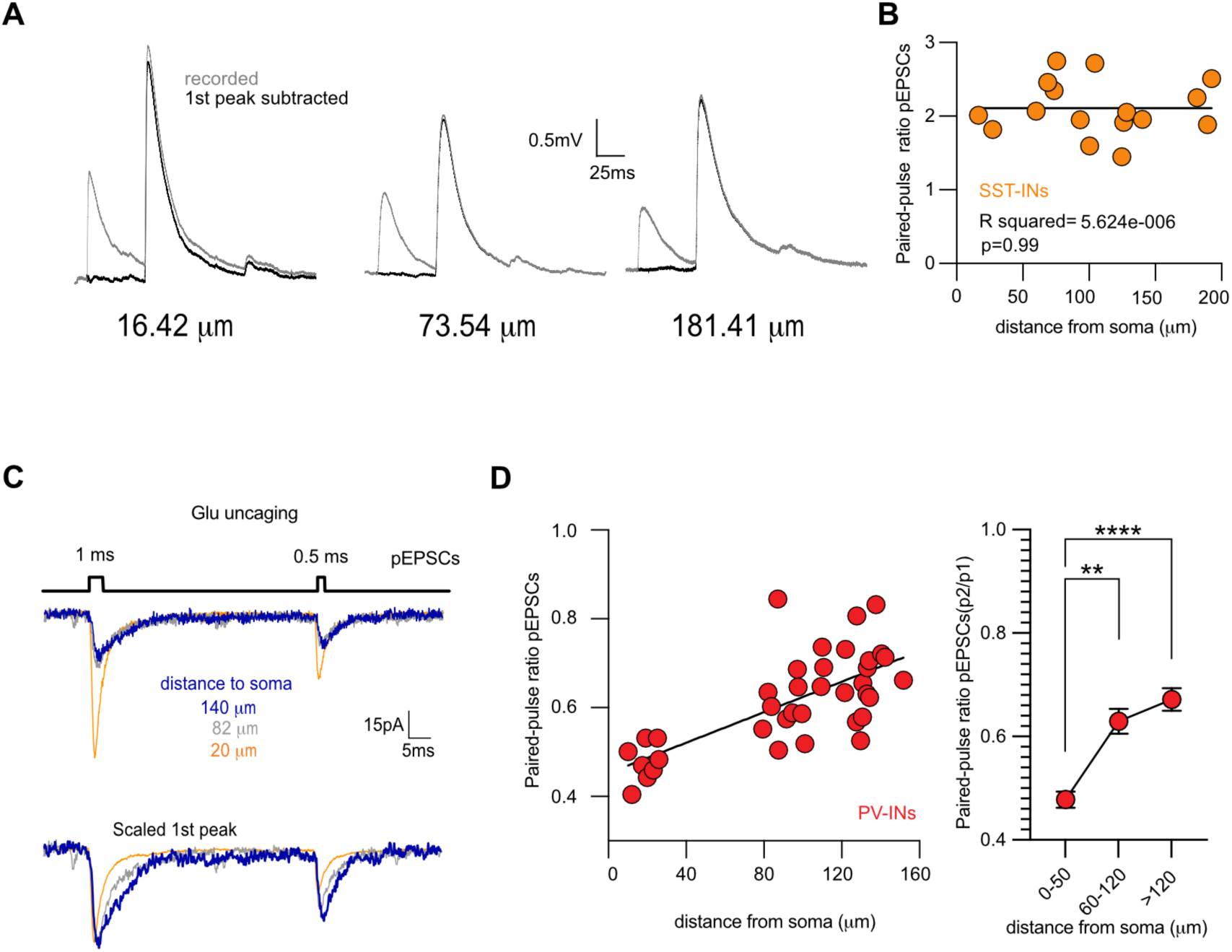
Integration of single and double quanta in SST+ dendrites in the current-clamp configuration and Double quanta integration in PV+ IN dendrites mimicking short-term depression. **A)** Example pEPSP traces of single and double quanta integration, separated by 50 ms (in gray), along an SST+ IN dendrite. In black the isolated second pEPSPs after subtraction of the first evoked response. **B)** Quantification of double quanta integration at different dendritic locations (n=16) expressed as paired-pulse ratio. In black the linear fit. C) *Top:* Double and single quanta integration mimicking a paired pulse depression at different dendritic locations (color coded). *Bottom:* Normalized traces scaled for the single quanta (2^nd^ peak). D) *(left)* Quantification of double quanta integration at different dendritic locations by using paired pulse ratio (n=37). In black the linear fit (p<0.0001 R^2^=0.49). *(right)* Histogram of binned PPR for values reported in B. Data are presented as mean±SEM (0-50: 0.48±0.01; 60-120 :0.63±0.02; >120: 0.67±0.02. 0-50 vs 60-120 p=0.002; 0-50 vs >120 p<0.0001, Kruskal Wallis test with Dunn’s correction.

**Figure S7:**
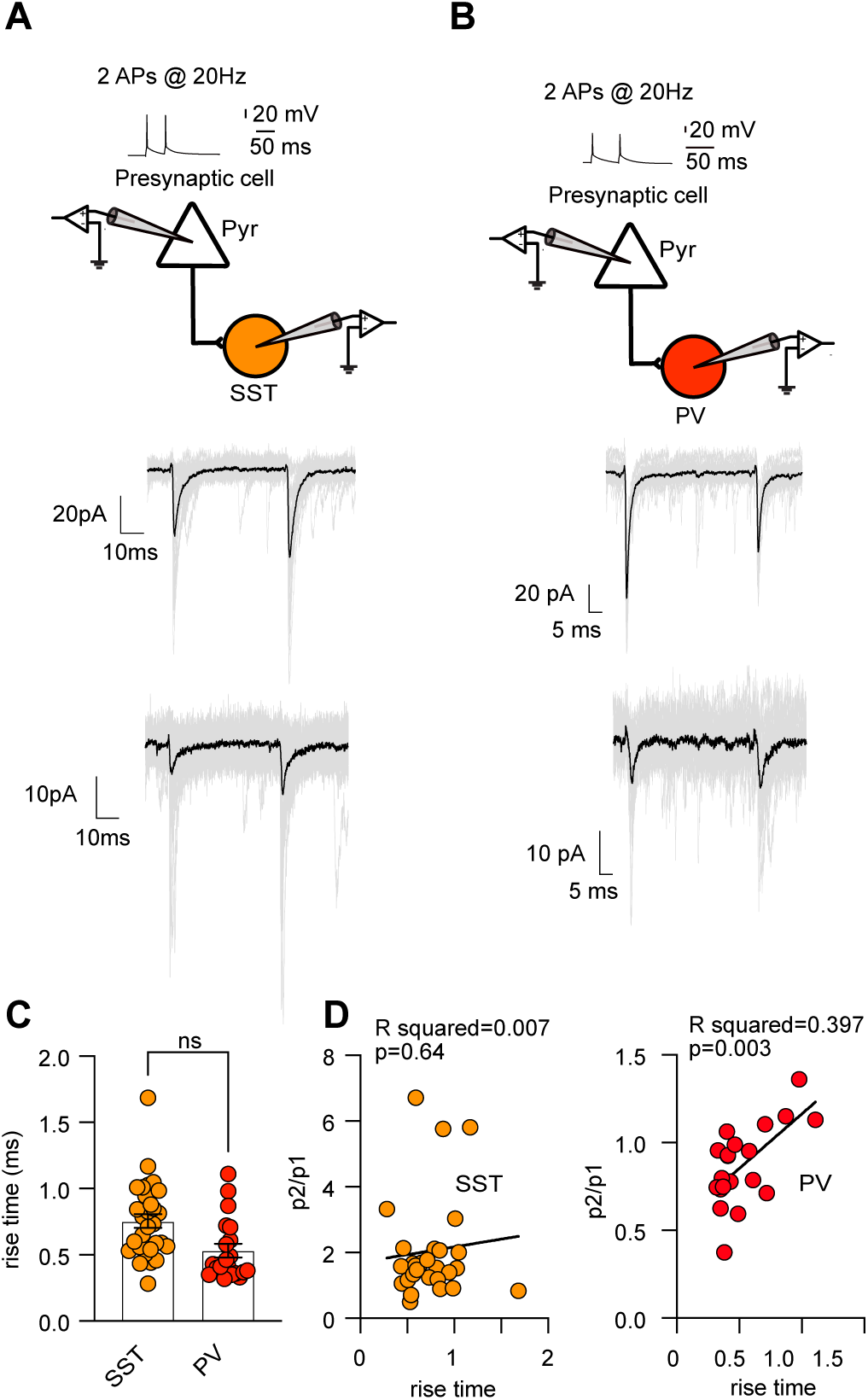
Paired recordings suggest a distance-dependent gradient of short-term plasticity in PV-INs but not SST-INs. **A)** *(top)* Illustration of dual whole-cell patch clamp recording used to probe unitary synaptic connections between L2/3 PNs and SST-INs. (*bottom*) Two example traces of EPSCs recorded in response to a paired-pulse stimulation in a single L2/3 PN. The two examples differ in rise time and amplitude. **B)** Same as in **A** but for unitary connections onto PV-INs. **C)** Summary data plot of EPSC rise time (first peak) recorded in SST(orange) and in PV(red) IN. Data are presented as mean±SEM (SST: 0.66±0.05 n=37; PV: 0.53±0.06 n=17, p=0.06 Mann Whitney test)**. D)** Summary plot of paired pulse ratios in function of rise time for the different recorded synaptically connected cell pairs. The black lines represent the linear fit (SST p=0.64, R^2^=0.007; PV p=0.03, R^2^=0.39.

**Figure S8:**
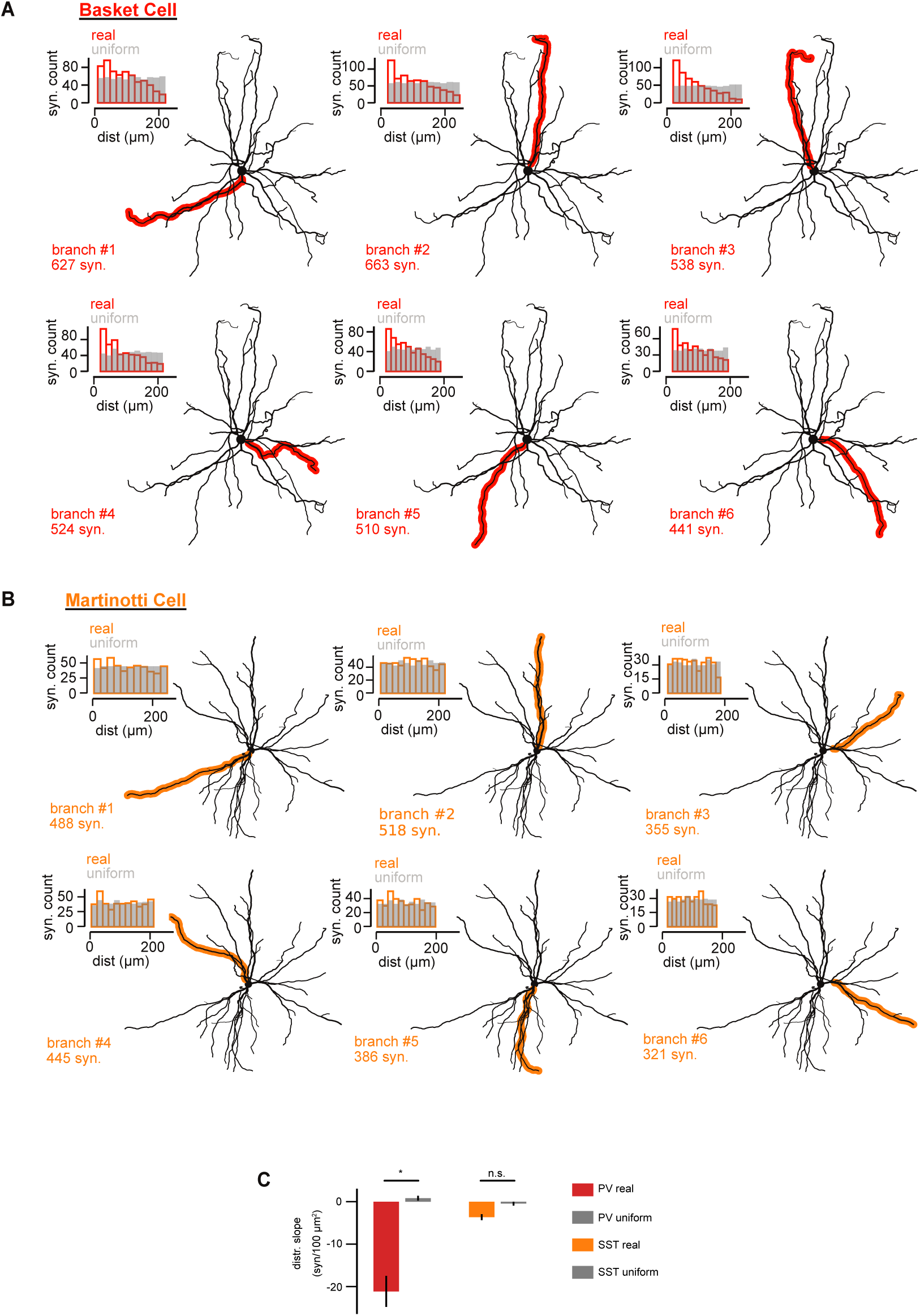
Dendritic branches used for numerical simulations in the PV-IN and SST-IN models. **A)** Morphology of a V1 basket cell reconstructed from serial electron microscopy imaging highlighting the six distinct dendritic branches used for numerical simulations of the PV-IN model. Histogram (top inset) of the real synapse distribution (colored) and its uniform-density surrogate (gray, see Methods) for the different individual dendritic branches. **B)** Same as **A** but for the Martinotti Cell used in the SST-IN model. **C)** Distributions slopes (of the synaptic count vs distance to soma) in the real synaptic distributions and uniform surrogates. Basket cell, real: -21.1±3.7 syn./100μm, uniform: 0.8±0.6 syn./100μm, n=6 branches, p=3e-2, Wilcoxon test. Martinotti cell, real: -3.6±0.7 syn./100μm, uniform: -0.5±0.5 syn./100μm, n=6 branches, p=6e-2, Wilcoxon test.

**Figure S9:**
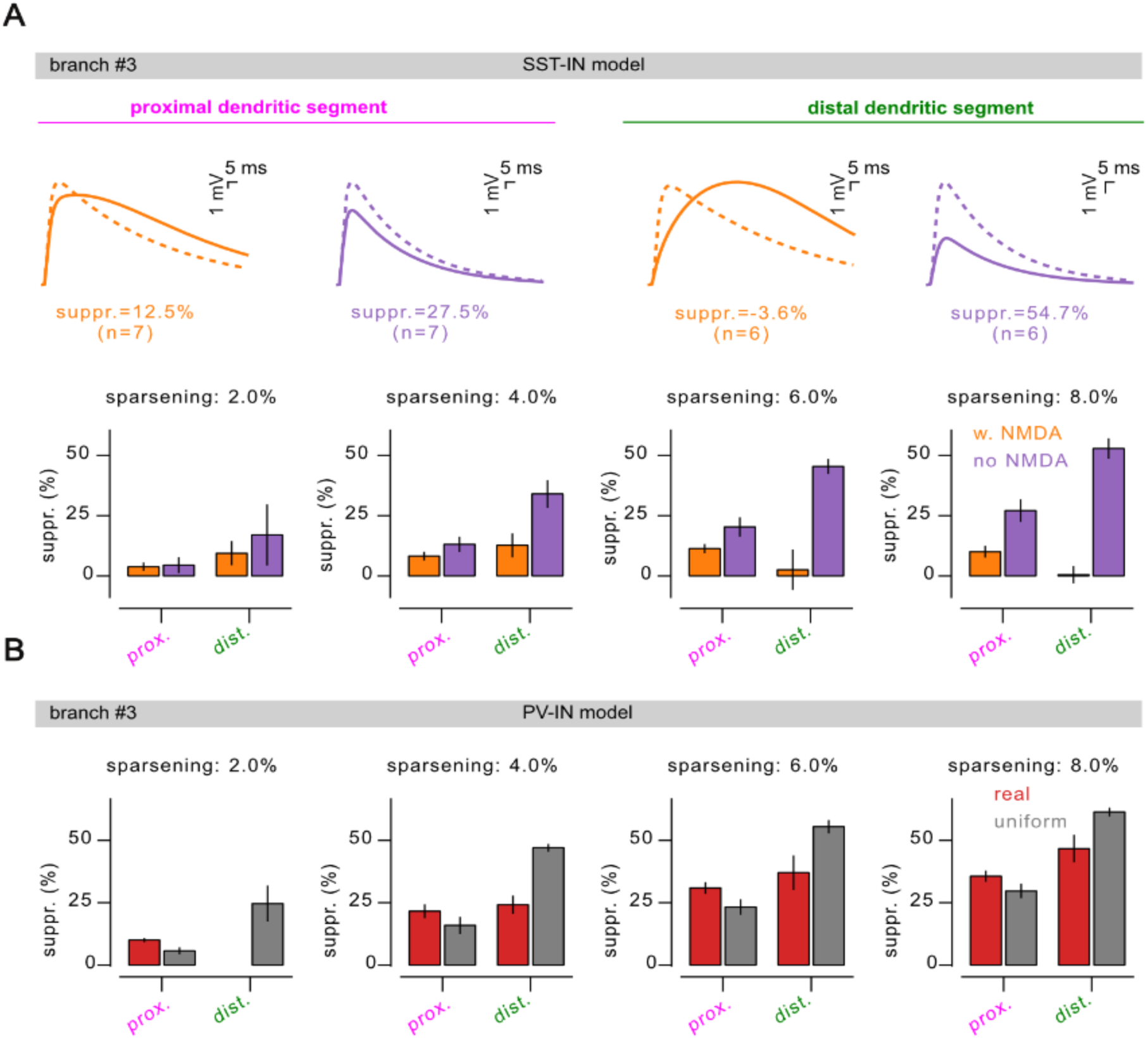
Distal and proximal integration upon increasing number of co-active inputs in dendritic segments of the morphologically-detailed interneuron models. Related to Fig. 4. We vary the sparsening variable that sets synaptic recruitment from 2% (left) to 8% (right) in the PV-model (**A**) and SST-model (**B**) with their respective control (see main text, “uniform” distribution for the PV-model and “no-NMDA” setting for the SST-model. Note the very high level of suppression (>50%) for sparsening above 4% in the PV-model. Note also the appearance of supra-linear integration (suppression <0%) in the distal segment of the SST-model in presence of NMDAR at a sparsening of 8%. **C)** Example V_m_ response in a branch in the SST-model at a sparsening of 8%. Integration from the distal segment stimulation exhibits supra-linear integration (suppression <0%).

**Figure S10:**
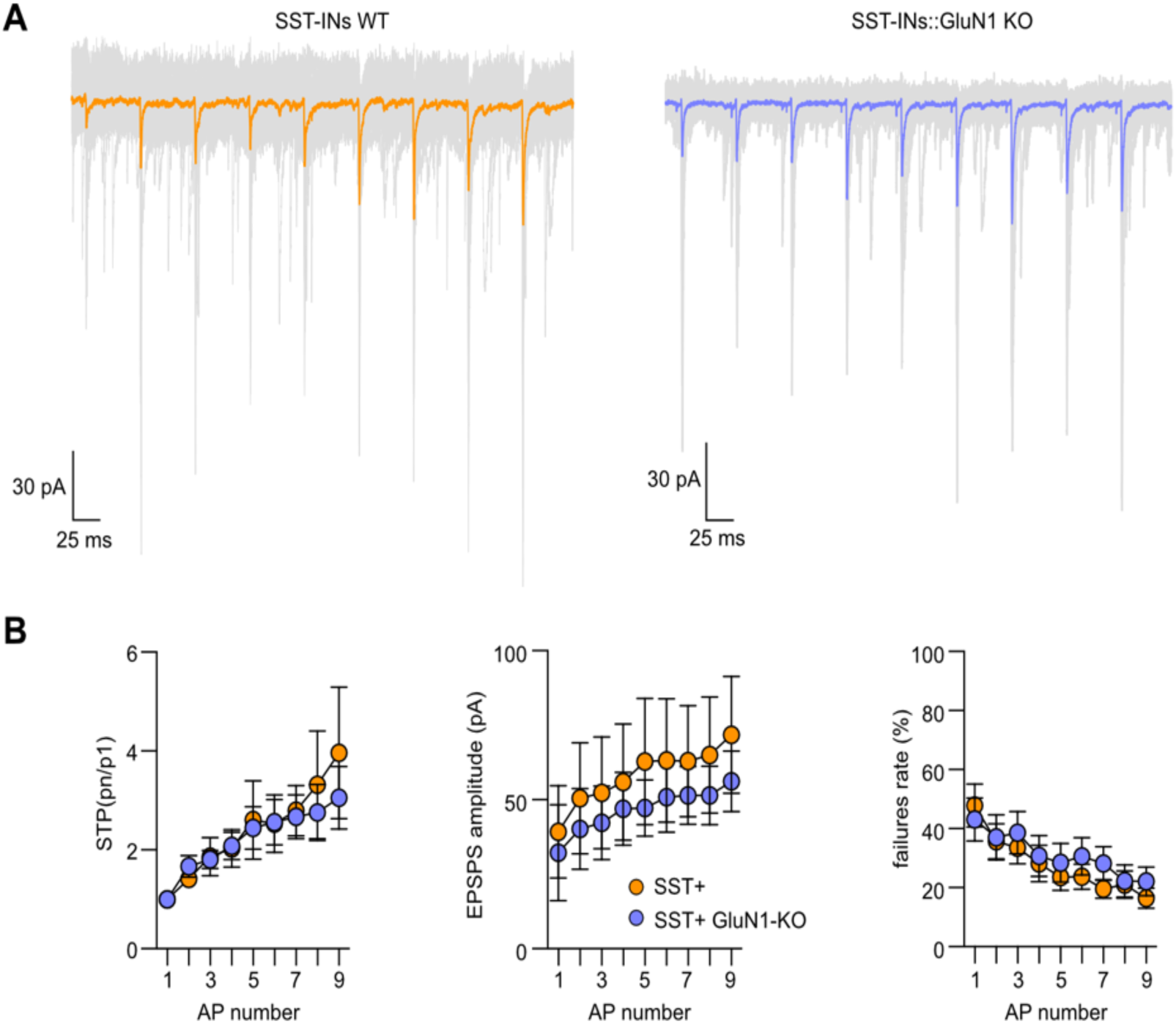
Synaptic transmission between L2/3 PN and L2/3 SST-INs is not altered in SST::GluN1 KO mice. **A)** Representative traces of uEPSCs recorded in a postsynaptic L2/3 SST-IN held at -70 mV in WT (left) and in SST-Cre::GluN1flox mice after a train of action potentials (9 @ 40 Hz) induction in a presynaptic L2/3 pyramidal neuron (PN). Single repetitions (30 sweeps) are in faint gray, average is in full color. **(B)** Summary plot of short-term plasticity, uEPSCs amplitude and failures rate (first EPSC of the train of stimulation) for the unitary connection between L2/3 PN and L2/3 SST-INs in WT (n = 14) and in SST-Cre::GluN1flox mice (n = 9).

## Supplementary Tables

**Supp. Table 1.** Passive properties of interneuron models.

| Supp. Table 1. Passive properties of interneuron models. |  |  |  |  |
| --- | --- | --- | --- | --- |
| Property | Soma | Proximal dendrite | Distal dendrite | Constraint/Reference |
| *PV model* |  |  |  |  |
| Leak conductance ($g_{pas}$) | $4.37 \cdot 10^{-4}$ S.cm<sup>-2</sup> | $4.37 \cdot 10^{-4}$ S.cm<sup>-2</sup> | $4.46 \cdot 10^{-5}$ S.cm<sup>-2</sup> | Adjusted to get a somatic input resistance of 100MΩ while keeping the prox./dist ratio of (11). |
| Resting Membrane Potential ($e_{pas}$) | -70 mV | -70 mV | -70 mV | (56, 57) |
| Membrane capacitance ($cm$) | 1.2 μF.cm<sup>-2</sup> | 1.2 μF.cm<sup>-2</sup> | 1.2 μF.cm<sup>-2</sup> | (58) |
| Axial Resistance ($Ra$) | 172 $\Omega$.cm | 142 $\Omega$.cm | 142 $\Omega$.cm | (58) |
| *SST model* |  |  |  |  |
| Leak conductance ($g_{pas}$) | $3.5 \cdot 10^{-5}$ S.cm<sup>-2</sup> | $3.5 \cdot 10^{-5}$ S.cm<sup>-2</sup> | $3.5 \cdot 10^{-5}$ S.cm<sup>-2</sup> | Adjusted to get a somatic input resistance in 300 +/- 40 MΩ (56), here 285MΩ |
| Resting Membrane Potential ($e_{pas}$) | -60 mV | -60 mV | -60 mV | (60, 61) |
| Membrane capacitance ($cm$) | 1.2 μF.cm<sup>-2</sup> | 1.2 μF.cm<sup>-2</sup> | 1.2 μF.cm<sup>-2</sup> | (58) |
| Axial Resistance ($Ra$) | 172 $\Omega$.cm | 142 $\Omega$.cm | 142 $\Omega$.cm | (58) |

**Supp. Table 2.**
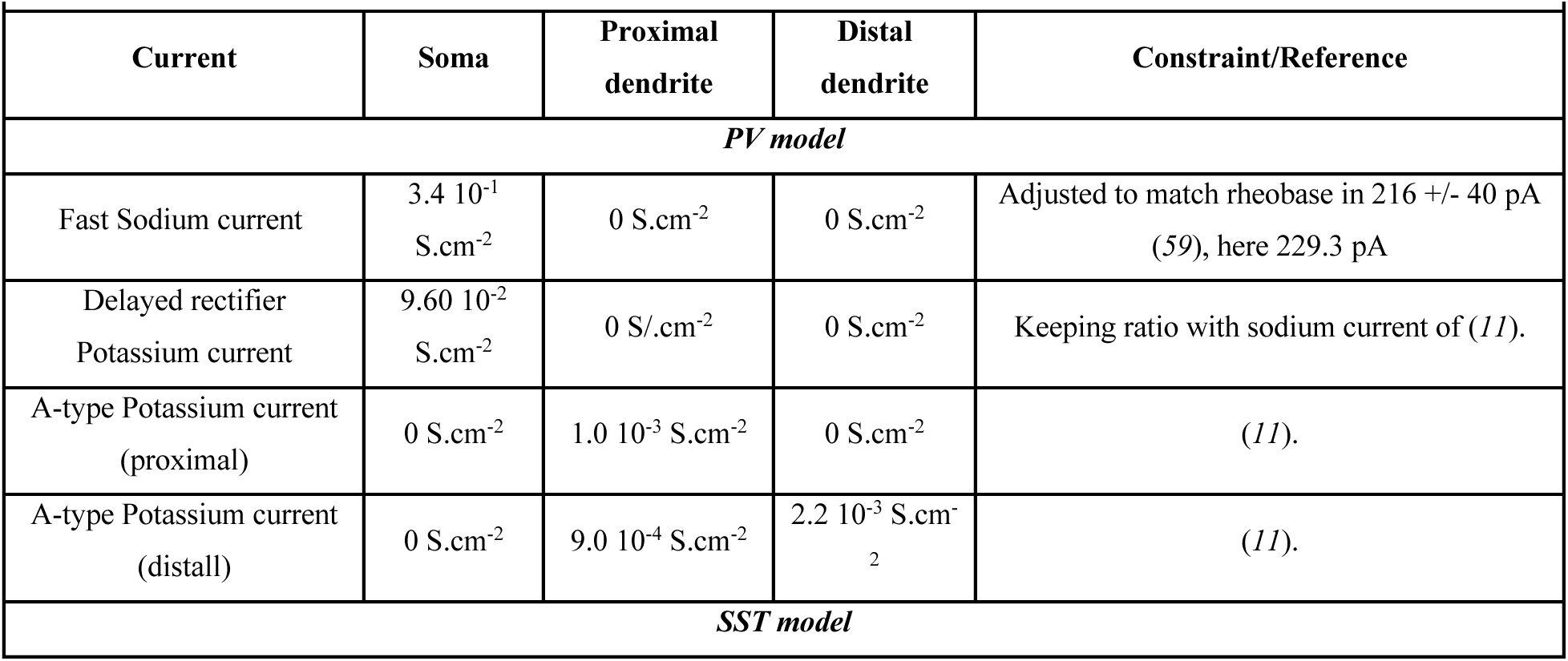
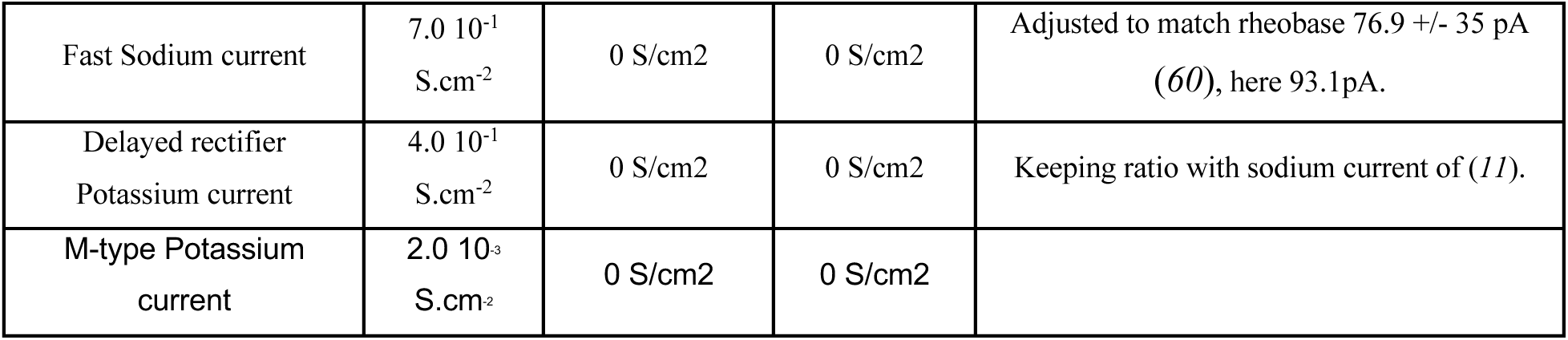
Active current densities in the interneuron models.

**Supp. Table 3.** Synaptic properties of the interneuron models.

| Supp. Table 3. Synaptic properties of the interneuron models. |  |  |  |
| --- | --- | --- | --- |
| Receptor | Quantal | Decay | Constraint/Reference |
| *PV model* |  |  |  |
| AMPA | 0.8 nS | 2 ms | (61) |
| GABA | 2 nS | 5.5 ms | (61) |
| *SST model* |  |  |  |
| AMPA | 0.8 nS | 2 ms | (61) |
| NMDA | 1.2 nS | 80 ms | (61)<br>Constrained from NMDA/AMPA ratio=1.5 |
| GABA | 2 nS | 5.5 ms | (61) |

## Methods

### Mice

Animals (C57BL/6 background) were housed in the Paris Brain Institute animal facility accredited by the French Ministry of Agriculture for performing experiments on live rodents under normal light/dark cycles. Work on animals was performed in compliance with French and European regulations on care and protection of laboratory animals (EC Directive 2010/63, French Law 2013–118, February 6th, 2013). All experiments were approved by local the Ethics Committee #005 and by French Ministry of Research and Innovation. Experimental data was obtained from adult (P25-P70) mice. Both male and female mice were used with the following genotypes: SST-IRES-Cre (SSTtm2.1(cre)Zjh/J; JAX 013044) X Ai9 (Gt(ROSA)26Sortm9(CAG-tdTomato); JAX 007909); PV-Cre (Pvalbtm1(cre)Arbr/J; JAX 008069) X Ai9 (Gt(ROSA)26Sortm9(CAG-tdTomato); SST-IRES-Cre (SSTtm2.1(cre)Zjh/J; JAX 013044) X PSD-95-ENABLED (B6;129-Dlg4tm1.1Hnz/J; JAX 026092); PV-Cre (Pvalbtm1(cre)Arbr/J; JAX 008069) X PSD-95-ENABLED (B6;129-Dlg4tm1.1Hnz/J; JAX 026092). Animals were maintained on a 12-h light/dark cycles with food and water provided ad libitum.

### Slice preparation

Acute parasagittal slices (320 μm) were prepared from adult mice, aged between postnatal days 25 and 70. Mice were deeply anesthetized with a mix of ketamine/xylazine (mix of i.p. ketamine 100 mg/kg and xylazine 13 mg/kg) and perfused transcardially with ice-cold cutting solution containing the following (in mM): 220 sucrose, 11 glucose, 2.5 KCl,1.25 NaH_2_PO_4_, 25 NaHCO_3_, 7 MgSO_4_, 0.5 CaCl_2_. After the perfusion the brain was quickly removed and slices prepared using a vibratome (Leica VT1200S). Slices containing S1 barrel field were transferred to ACSF solution at 34°C containing (in mM): 125 NaCl, 2.5 KCl, 2 CaCl_2_, 1 MgCl_2_, 1.25 NaH_2_PO_4_, 25 NaHCO_3_, 15 Glucose for 15–20 min. After the period of recovery, slices were kept at room temperature for a period of 5-6 h.

### Electrophysiology

Whole-cell patch-clamp recordings were performed close to physiological temperature (32–34°C) using a Multiclamp 700B amplifier (Molecular Devices) and fire-polished thick-walled glass patch electrodes (1.85 mm OD, 0.84 mm ID, World Precision Instruments); 3.5–5 MOhm tip resistance. For voltage-clamp recordings, cells were whole-cell patched using following intracellular solution (in mM): 90 Cs-MeSO_3_, 10 EGTA, 40 HEPES, 4 MgCl_2_, 5 QX-314, 2.5 CaCl_2_,10 Na_2_Phosphocreatine, 0.3 MgGTP, 4 Na_2_ATP (300 mOsm pH adjusted to 7.3 using CsOH). Extracellular synaptic stimulation was achieved by applying voltage pulses (20 μs, 5–50 V; Digitimer Ltd, UK) via a second patch pipette filled with ACSF and placed 20–40 μm from soma. For current clamp glutamate uncaging experiments patch pipettes were filled with the following intracellular solution (in mM): 135 K-gluconate, 5 KCl, 10 HEPES, 0.01 EGTA, 10 Na_2_phosphocreatine, 4 MgATP, 0.3 NaGTP (295 mOsm, pH adjusted to 7.3 using KOH). The membrane potential (Vm) was held at -60 mV, if necessary, using small current injection (typically in a range between -50 pA and 200 pA). Recordings were not corrected for liquid junction potential. Series resistance was compensated online by balancing the bridge and compensating pipette capacitance. For glutamate uncaging experiments Alexa 594 (20 μM) was added to the intracellular solution daily. In all experiments data were discarded if series resistance, measured with a -10 mV pulse in voltage clamp configuration, was >20 MΩ or changed by more than 20% across the course of an experiment. For current clamp experiment cells were excluded if input resistance varied by more than 25% of the initial value. AMPA-EPSCs were recorded at -70 mV in the presence of the GABAA blocker picrotoxin (100 μM, Abcam). To record NMDA-EPSCs, NBQX (10 μM, Tocris) was added to the ASCF and the resting membrane potential changed to +40 mV. All recordings were low-pass filtered at 10 kHz and digitized at 100 kHz using an analog-to-digital converter (model NI USB 6259, National Instruments, Austin, TX, USA) and acquired with Nclamp software (*62*) running in Igor PRO (Wavemetrics, Lake Oswego, OR, USA). For all the experiments L2/3-PV+-INs and L2/3-SST+-INs were identified using PV-Cre and SST-cre mice crossed with the reporter mouse line (Ai9, (ROSA)26Sortm9 (CAG-tdTomato)Hze/Jt; JAX 007909). The intracellular solution used for the experiments in which a presynaptic cell was depolarized by a train of action potentials to induce neurotransmitter release on a post-synaptic cell/element was (in mM): 130 K-MeSO3, 4 MgCl2,10 HEPES, 0.01 EGTA, 4 Na2ATP, 0.3 NaGTP (300 mOsm, pH 7.3 adjusted with NaOH). Paired recordings connections were probed using a train of 5/9 action potentials at frequency of 20 Hz. APs were initiated by brief current injection ranging from 1000–2000 pA and 1–5 ms duration. After a clear identification of synaptic connection (mean of 30 sweeps with clear AMPA current), post-synaptic cell membrane potential was brought from the initial -70 mV membrane potential to +40 mV to record the NMDAR-EPSCs in the presence of blockers as described before. To visualize single boutons in the experiments targeting SFiGluSnFR.A184V positive dendritic elements, we added 40 μM Alexa 594 in the intracellular solution.

### Transmitted light and fluorescence imaging

Neurons were visualized using infrared Dodt contrast (Luigs and Neumann, Ratingen, Germany) and a frame transfer CCD camera (Infinity-Lumenera). These components were mounted on an Ultima two-photon laser scanning head (Bruker, USA) based on an Olympus BX61W1 microscope, equipped with a water-immersion objective (60x, 1.1 numerical aperture, Olympus Optical, Japan). The somas of somatostatin (SST) and parvalbumin (PV) interneurons were identified using a td-Tomato reporter mouse line (see animals) combined with fluorescence imaging. Two-photon excitation was performed with a pulsed Ti:Sapphire laser (DeepSee, Spectra-Physics, France) tuned to 810 or 840 nm for imaging neuronal morphology. Individual neurons were patch-loaded with Alexa 594 (20 μM). In some instances, we used a transmitted light PMT mounted after the Dodt tube to acquire a laser-illuminated contrast image simultaneously with the 2PLSM image. This dual imaging mode was used to position stimulation electrodes of sucrose puffing pipettes close to a spatially isolated dendrite, identified from maximal intensity projections of 2PLSM images. In addition to the light collected through the objective, the transmitted infrared light was collected through a 1.4 NA oil-immersion condenser (Olympus), and reflected on a set of substage photomultiplier tubes (PMTs). SFiGluSnFR.A184V fluorescence were filtered using hq525/70 nm bandpass filters (Chroma) and detected with gallium arsenide phosphate-based PMTs (H10770PA-40, Hamamatsu Photonics), and Alexa Fluor 594 fluorescence were filtered using hq607/45 bandpass filters (Chroma) and detected using gallium arsenide phosphate-based PMTs (H10770PA-40, Hamamatsu Photonics). 200 nm yellow-green fluorescent beads (Life Technologies) were used to estimate the point spread function (PSF) of the microscope system. The measured PSF at 810 nm had lateral dimensions of 368±5.19 nm (full width at half maximum, n =20).

### Stereotaxic viral injections

Adult mice were bilaterally injected in the primary somatosensory cortex barrel field (S1-BF) by or unilaterally in the primary visual cortex (V1) using a 10 μl Hamilton syringe (1701RN; Phymep, France) with a borosilicate glass capillary (O.D. 1.0mm; I.D.0.50mm; 10 cm length item #: BF100-50-10 WPI) pulled and beveled mounted on automatic injector used to infuse between 100 and 450 nl of virus at a rate of 60 nl/min using the following coordinates relative to Bregma: S1 anterior-posterior [AP], -0.5mm; medial lateral [ML], ± 3.2 mm and dorso ventral [DL] -0.2 mm relative to brain surface; V1(anterior-posterior [AP], -4.1mm; medial lateral [ML], ± 2.8 mm and the dorso ventral [DL], -0.2 mm relative to brain surface. Following the injection, the beveled glass capillar was left in place for 4 minutes to allow full diffusion of the viruses. E*x vivo* electrophysiology experiments were performed at least 2/3 weeks after the surgery. For monitoring glutamate release at individual glutamatergic synapses, PV- or SST-cre mice were stereotaxically injected with AAV9.CAG-FLEX.SFiGluSnFR.A184V (Addgene plasmid #106205; (*39*)). For in vivo two photon calcium imaging mice were injected with AAV9 Syn-Flex-GCaMP6s-WPRE-SV40 (2.5x1013 AddGene-100845). To trace dendritic elements from SST or PV positive neurons in the STED experiment we inject AAV1.Flex.tdTomato (Addgene-28360) in S1-BF.

### Estimation quantal miniature EPSCs

The morphology of L2/3 SST- and PV-INs was visualized using fluorescence imaging of patch-loaded Alexa 594 (20 μM). The extracellular solution was modified to contain in mM: 0 CaCl2 and 3 MgCl2 in the presence of TTX 0.5 μM (*63*). Neurons were patch loaded with Alexa 594 for at least 20 min to visualize cell morphology using two-photon microscopy with a laser tuned to 820 nm and modulated using a Pockels cell (Conoptics, Danbury, CT). This strategy allowed us to identify the proximal and distal dendrites of the patched neuron. A puff pipette was then placed at 20-30 μm from the selected dendrite where an extracellular hypertonic sucrose solution (350 mOsm) was puffed for 2 seconds at 0.40 bar positive pressure. Single evoked miniature events were then isolated from the 2 second sucrose puffing period recorded in 20 sweeps. On average 120 events were used for analysis. For single experiments rise time estimation was performed by fitting single events with a triexponential function. Fitted events have been subsequently aligned for their respective 10% rise time which finally resulted in the reported amplitude and kinetics values (fig S3). To reduce possible contamination of distally recorded events by spontaneous proximal events in PV-INs due to their elevated basal rate of mEPSCs we used an exclusion criteria based on rise times. During estimation of sucrose evoked mEPSCs in distal dendritic regions in PV-INs we excluded events whose rise time was within mean+2SD the rise time of the sucrose mEPSCs obtained on the same cell. This criteria allowed us to remove spontaneous proximal events that occurred during the sucrose puffing period. The distance used for proximal and distal evoked events was respectively 13.55 ± 3.46 and 113.6 ± 4.89 nm for PV-INs and 22.51 ± 2.60 and 124.2 ± 9.61 nm for SST-INs.

### Glutamate uncaging

L2/3 PN and L2/3 INs morphology was visualized using fluorescence imaging of patch-loaded Alexa 594 (20 μM). The output of two pulsed Ti:Sapphire (DeepSee, Spectra Physics) lasers were independently modulated to combine uncaging of MNI-glutamate and Alexa 594 imaging. The imaging laser beam was tuned to 820 nm and modulated using a Pockels cell (Conoptics, Danbury, CT). For uncaging, the intensity and duration (0.5s or 1s for single and double quanta experiment) of illumination of the second Ti:Sapphire laser tuned to 720 nm was modulated using an acousto-optic modulator (AA Opto-Electronic, France). MNI-caged-L-glutamate (10 mM) was dissolved in a solution containing (in mM): NaCl 125, glucose 25, KCl 2.5, HEPES 10, CaCl_2_ 2 and MgCl_2_ 1 (pH 7.3) and constantly puffed using a glass pipette (res 1.5–2 MOhm) placed in the proximity of a selected dendrite. Multiple uncaging locations (6–12) were placed adjacent to visually identified spines for PNs or in proximity to the shafts for INs (1 μm) and not closer to each other than 2 μm to reduce possible effect of glutamate spillover between locations. For the uncaging experiments in INs the uncaging spots were placed along the dendritic shaft at the level of synaptic “hot-spots” identified by single events rise time as described in (*13*, *64*). Laser intensity was adjusted to obtain individual pEPSP of a variable amplitude. The arithmetic sum (always corrected for not perfect simultaneous activation) was used to compare the obtained values (expected) versus the recorded EPSP elicited by an increasing number of inputs recruited by glutamate uncaging. An input-output curve was obtained by plotting the amplitude of the expected EPSP versus the amplitude of recorded EPSP. Non-linearity was calculated as previously described (*65*). Different inter event time intervals were tested during the quasi simultaneous activation of inputs: 0.12 and 1, 5 or 8 ms. In a subset of experiments laser power used to evoke glutamate uncaging was adjusted in order to obtain pESPCs with similar amplitudes than the mean amplitude of the quantal miniature EPSCs recorded during sucrose puff at proximal and distal locations. This initial calibration was performed in the voltage clamp configuration. Amplitude of quantal adjusted pEPSPs (qEPSPs) was also obtained in a separate group of experiments by shifting the recording conditions to the current clamp configuration while maintaining uncaging location and laser intensity. This allowed us to obtain the following values: qEPSCs PV-INs proximal= 32.90 ± 1.27 pA, qEPSPs PV-INs proximal 0.73 ± 0.06 mV; qEPSCs PV-INs distal= 16.40 ± 1.39 pA, qEPSPs PV-INs distal 0.47 ± 0.06 mV; qEPSCs SST-INs distal= 17.06 ± 0.17 pA, qEPSPs SST-INs distal 0.83 ± 0.17 mV. The drugs used for uncaging experiments were placed in the puff pipette and in the bath. In glutamate uncaging experiments involving specific receptors antagonists and/or channel blockers, the drugs were present in both the puff pipette solution and the bath solution. The concentration of NMDA blockers, D-AP5 and MK-801, used in the puff pipette solution was 500 μM and 50 μM respectively following the conditions described in (*66*). In the bath solution concentration of D-AP5 and MK-801 used was 50 μM and 20 μM respectively. In a subset of experiments we also blocked voltage gated calcium and sodium channels in addition to NMDARs (described as “blockers cocktail). For that TTX 1μM, NiCl 100μM, Nifedipine 30μM were added to the puff and bath solution.

### iGluSnFR Imaging

Acute brain slices from animals previously injected with AAV9.CAG-FLEX.SFiGluSnFR.A184V were performed as described above. L2/3 PNs were subsequently patch loaded with Alexa 594 (40 μM) and the axon visualized using two-photon imaging. Axonal boutons were easily identified after 30-40 min of intracellular loading in whole cell patch-clamp configuration. Using simultaneous two-photon imaging of Alexa 594 and iGluSnFR fluorescence signals, putative points of contact were first identified at 820nm. The genetically encoded glutamate sensor, iGluSnFR-A184V was selectively expressed in SST- or PV-INs. Presence of positive synaptic connections was probed by placing a linescan (∼7 μm at 0.76 ms per line) over the identified potential synaptic connection and variation of iGluSnFR fluorescence signals (imaged at 920 nm) monitored in response to evoked action potentials via current injection in the presynaptic L2/3 PN. Trains of 7 action potentials (@10 Hz) repeated at a frequency 0.33Hz were used to imaged glutamate release into PV- and SST-INs. Variations in iGluSnFR fluorescence were expressed as ΔF/F time series traces constructed from linescan images. The fluorescence as a function of time was averaged over visually identified pixels corresponding to width of the presynaptic bouton (identified using Alexa 594 fluorescence and overlapping with the iGluSnFR expressing dendrite) and then averaged over individual trials, resulting in a single fluorescence trace as a function of time (F_dendrite_(t)). The background fluorescence (F_back_(t)) was estimated similarly (identical spatial line length), but from a location not on a labeled structure. This F_back_(t) trace was then fit with a single exponential function and then subtracted from F_dendrite_(t) in order to correct for bleaching. The background corrected traces (F(t)) was then converted to the final ΔF/F(t) according to Equation 1:

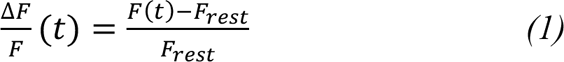

where F_rest_ is the time averaged (100 ms window) fluorescence before stimulation and F(t) is the time-dependent fluorescence transient

### Optical fluctuation analysis

The estimation of number release sites (N_max_ves) and probability of release (P_Ves_) from single boutons using iGluSnFR fluorescence signals was performed using fluctuation analysis inspired in the method previously described for estimating number and open probability of voltage gated calcium channels (VGCCs; (*36*, *37*)). We considered simplistically that if neurotransmitter release at single point of contact is governed by binomial statistics, then the coefficient of variation of the dark/shot noise subtracted iGluSnFR fluorescence variance is given by:

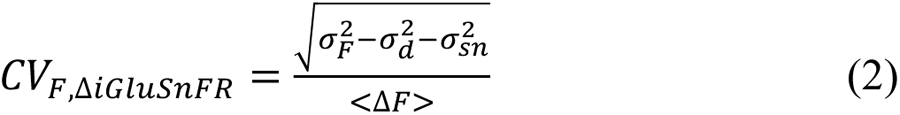

where 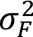 is the total signal variance, 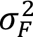 is the dark noise and 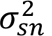 represents the shot noise contributions, where the shot noise variance was calculated by imaging a pipette filled with Alexa 488 with varying laser intensities. To ensure that variance was related to fluctuations in neurotransmitter release and to minimize other sources of variance, including optical drift we corrected the variance with a trial to trial correction for drift as reported in (*67*) by using the equation:

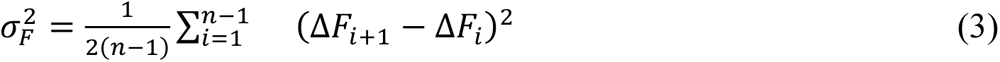

where n represents the number of trials. This normalization minimizes the variance increase given by a run up or run down of fluorescence during the recording. Recordings in which the baseline variance (calculated from a 100 ms window before AP induction) was larger than the 99% confidence interval of the shot noise expected variance were not included in the analysis. Failure rate estimation was defined using a thresholding approach on the F(t) trace. We systematically tested multiple different thresholds. For that we compared the mean F_iGluSnFR_ value (calculated from a 10 ms window from the F(t) trace just after presynaptic evoked action potentials) to the mean + 0.5, 1 or 2 times the SD of the baseline (prior action potential). Release successes were defined if F value after AP was larger than threshold. In order to optimally define the best threshold for each individual experiment we computed variance from the failure trials defined for the different thresholds and compared to predicted shot noise variance. All successes were further analyzed as ΔF/F. To estimate the amplitude of release successes, we fit averaged ΔF/F traces with the following equation (*68*), using a least-square algorithm implemented in IgorPro (Wavemetrics):

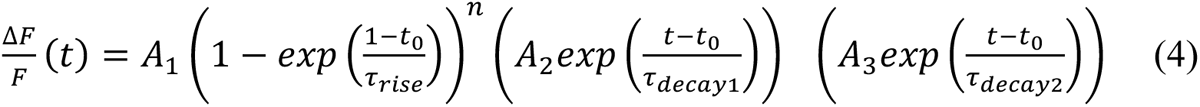

Assuming that neurotransmitter release at single point of contact is governed by binomial statistics the probability that an action potential fails to release any synaptic vesicle is given by the equation:

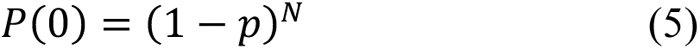

where p is the release probability of individual vesicles and N is the number of release sites. Equation 5 together with CV_F_,_iGluSnFR_ can be used to solve for N and p.

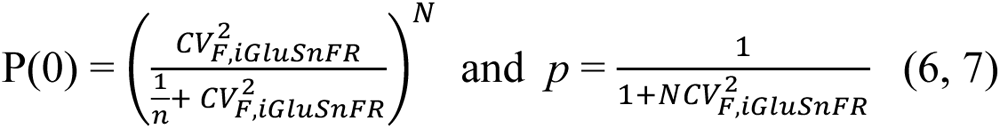

### PSD95 labelling in cortical interneurons

We examined the distribution of excitatory synaptic inputs along the somatodendritic compartment in SST- and PV-INs using transgenic mouses line that conditionally express Venus-tagged PSD95 under the control of the SST- or PV- promoter (PSD95-Enabled (*42*) × SST Cre or PV-Cre respectively. Stereotaxic viral injections of AAV1.Flex.tdTomato was used to visualize interneuron morphology in such transgenic mouse lines.Two/three weeks after stereotaxic viral infections, SST-PSD-95-CreNABLED and PV-PSD-95-CreNABLED adult mice were deeply anesthetized with a mix of ketamine/xylazine (mix of i.p. ketamine 100 mg/kg and xylazine 13 mg/kg) and perfused transcardially with ice-cold PBS 1X (ET330-A) followed by ice-cold PFA 4% freshly prepared. At least 30 ml of PFA was used per animal. After extraction, brains were immersed in PFA 4% for and additional 2 hours, at 4C, and sliced on the same day as the perfusion. Parasagittal slices (100 um) containing S1 were prepared at 4°C using a vibratome (Leica VT1200S). Slices were kept at 4°C in a PBS-Azide 0.4% solution. In order to amplify the endogenous PSD95-mVenus fluorescence signal brain sections were immunostained with an anti-GFP Nanobody coupled to ATTO 647N (GBA647N-100, ChromoTek). For that brain sections were washed for 10 minutes in Phosphate Buffer (PB) 0.1M, followed by three 10 min incubation in Tris Buffer Saline (TBS). Sections were subsequently incubated for 1 hour in a blocking solution composed of 10% Normal Goat Serum (NGS), 0.2% triton 100-X in TBS. Directly after the blocking solution, brain sections were incubated for 2 hours in a solution of 2% NGS, 0.2% triton 100-X, GFP-Booster ATTO647N (GBA647N-100, ChromoTek) ((1:2000) in TBS. Sections were then washed (3 X 10 minutes) in TBS and rinsed in PB 0.1M and mounted on microscope slides using ProLongGlass Antifade Mountant (ProLong Glass, Invitrogen P36982). Fluorescence images were acquired at least 3 days after mouting to allow mouting medium to cure. Brain slices were imaged with a Stimulated Emission Depletion (STED) fluorescence microscope (expert line - Abberior Instruments) using an Olympus 100X/1.4 NA oil objective lens and 775 nm STED Laser line. Excitation laser lines were at 561nm and at 640nm respectively for TdTomato and Atto 647N. STED Images were analyzed using Fiji.58.

### Surgery for in vivo 2P imaging

Mice were initially anesthetized with Isoflurane. Induction of anaesthesia was performed at 4% isoflurane, Iso-Vet, and an air flow of 250ml/min. Mice were then placed in a stereotactic device (Kopf) kept at 37 °C using a regulated heating blanket and a thermal probe and maintained under anaesthesia using 1-2% Isoflurane and an air flow of 250 ml/min until the end of the surgery. For calcium imaging, cranial window implantation and AAV viral stereotaxic injection was performed in the same surgical procedure. In order to prevent brain swelling and post-operative pain, anti-inflammatory (Dexamethasone, 2 µg/g Dexadreson, MSD) and analgesic drugs (Buprenorphine, 0.1 mg/kg Buprecare) were injected subcutaneously. After shaving the scalp, the skin was cleaned by wiping multiple times with an antiseptic solution and 70% alcohol. After injecting a local anaesthetic under the scalp (lidocaine, 10mg/kg -Xylovet or Laocaine) a section of the skin was removed using surgical scissors. The periosteum was then carefully removed and the skull was scrapped using a dental drill, in order to remove all residues. The surface of the skull was subsequently thoroughly cleaned and dried using cotton swabs and a physiological saline solution. Using the stereotactic device, the center of the cranial window was marked and a circular piece of bone was removed using a 3 mm biopsy punch (LCH-PUK-30 Kai Medical). This step of the surgical procedure was considered critical since damage to the dura would result in rapidly opacifying cranial windows. Once the brain was exposed, the AAV viral vectors carrying the genetically encoded calcium indicators (GCaMP6s) were stereotactically injected. For calcium imaging of SST-INs, AAV9 Syn-Flex-GCaMP6s-WPRE-SV40 (2.5x1013 AddGene-100845) was used respectively. Using a 10uL Gastight syringe 1701N (Hamilton), 200 nL of virus were injected in the visual cortex (from bregma: 3.0 RC; 2.4 ML) at a rate of 1 nL/s followed by a 5 minutes waiting period to prevent backflow. A round of 3mm glass coverslip (CS-3R Warner Instruments) was then placed over the craniotomy and glued in place using dental cement (Superbond). A stainless-steel head post (Luigs & Neumann) was then attached to the skull also using dental cement. For 3 days following the cranial windows surgery, mice were injected once a day with a mix of anti-inflammatory (Dexamethasone, 2 µg/g Dexadreson, MSD) and analgesic drugs (Buprenorphine, 0.1 mg/kg Buprecare). One week after the surgery, animals were transferred into an inverted light-dark cycle housing. Two weeks after surgery, mice were first habituated to the experimenter by handling and the following days, mice were head-fixed in the acquisition setup for increasing periods of time (5 – 45 minutes). The habituation of animals to the experimental setup lasted at least 5 days with a minimum of 5 sessions. At the end of the habituation period animals spontaneously transitioned between rest and running periods in the circular treadmill. Animals were imaged between Zeitgeber time 12 and 24. Imaging sessions lasted between 30 to 60 min and animals were imaged between 3-5 days. The location of the injection site in V1 was confirmed with intrinsic imaging experiments (*69*) as performed in our previous work (*70*).

### In vivo 2P Ca^2+^ imaging

In vivo two-photon calcium imaging was performed with an Ultima IV two-photon laser-scanning microscope system (Bruker), using a 20X, 1.0 N.A. water immersion objective (Olympus) with the femtosecond laser (MaiTai DeepSee, Spectra Physics) tuned to 920 nm for imaging of cells expressing GCaMP6s. Fluorescence light was separated from the excitation path through a long pass dichroic (660dcxr; Chroma, USA), split into green and red channels with a second long pass dichroic (575dcxr; Chroma, USA), and cleaned up with band pass filters (hq525/70 and hq607/45; Chroma, USA). Fluorescence was detected using both proximal epifluorescence photomultiplier tubes (gallium arsenide phosphide, H7422PA-40 SEL, Hamamatsu). Time-series movies of neuronal populations expressing GCaMP6s were acquired at the frame rate of 30Hz (512 × 512 pixels field of view; 1.13 μm/pixel). During the recording periods animals were free to run on a circular treadmill covered with a soft foam. This foam softened the movement of the animals and provided better traction (this was important since mice adapted more rapidly to the experimental setup and movements were more fluid during the recordings).

### Visual stimulation

Visual stimuli were presented at 60Hz on a Dell-2020 LED screen and synchronized to imaging and behavioral data via a photodiode (placed on the top right of the screen) that recorded the timing of each stimulus frame. Visual stimulation was controlled via the Psychopy software (*71*), version 2021.1.1. Full-field static gratings were presented for 2s (interleaved with 4s interstim intervals) with 2 contrast levels (half-contrast and full contrast), 8 different orientations evenly spaced from 0° to 157.5° and a spatial frequency of 0.04 cycle/degree. For two-photon experiments using a resonant scanner, the power source of the LED backlight of the monitor was synchronized to the resonant scanner turnaround points (when data were not acquired) to minimize light leak from the monitor (*72*).

### Reduced model of dendritic integration

Adapting the work of (*48*), we studied dendritic integration by implementing cable theory in a simplified model of the dendritic arborization. The overall dendritic structure is schematized in **Fig. 1I**, the morphology is made of 4 generations of branches with length 50µm. At the end of each generation *b*, each branch divides into two branches of a smaller diameter D_b+1_=T·D_b_ where T is the tapering factor. The root diameter (D_1_) was set to 1.0μm to match our proximal measurements (**Fig. S3**) and tapering factor was set to T=0.7 so that the diameter reached D_3_=0.5μm in the [100,150]μm of our distal measurements (**Fig. S3**). An isopotential spherical compartment of radius 15μm was added to model the soma. The membrane leak conductance was adjusted to gL=2.5pS/μm^2^ to match the ∼100MΩ input resistance for PV+ cells while the other cable properties were fixed to classical values with the membrane capacitance set to c_m_ =1μF/cm^2^ and the intracellular resistivity to R_i_=150Ω.cm. Synaptic integration was tested by adding synaptic conductance events with an instantaneous rise of amplitude 1nS and an exponential decay of τ=5ms. The reversal potential of the associated synaptic current was set to E_rev_=0mV. This morphology and electrical properties were implemented in the brian2 simulator (*73*) and simulated with a time step dt=0.025ms and with a spatial discretization of n=20 steps per branch.

### Numerical simulations of morphologically-detailed PV- and SST-IN models

The morphologically-detailed models were implemented within the NEURON simulation environment (*74*), version 8.2.3. From the Electron-Microscopy dataset of (*45*), we took the detailed morphological reconstructions of the Basket (cell-ID: 864691135396580129) and Martinotti (cell-ID: 864691135571546917) example cells shown in **Fig. 3G** for the PV-IN and SST-IN models respectively. To focus on dendritic integration and for numerical simulation efficiency, axonal compartments were removed from the reconstructions and only the somatic and dendritic compartments were simulated. The passive properties of the models were constrained from experimental measurements, see **Table S1**. **Fig 4-D** relies on passive properties only. To investigate the spiking output properties following dendritic integration, we next added a minimal set of active currents in **Fig 4E-G** and **Fig. 5A-D** for each model (**Table S2**). The action potential currents were made of a fast voltage-dependent sodium channel (*gnafin*) and a delayed rectifier potassium channel (*gkdrin*) with densities adjusted to match the rheobase of experimentally recorded cells (**Table S2**). For the PV-IN model, we also added A-type potassium channels (**Table S2**) as this was shown to strongly participate in dendritic integration in PV-INs (*11*). For SST-INs, we also added M-type potassium channels in the soma (**Table S2**) as those shape the regular-spiking response characterizing SST-INs spiking profile. Action potentials were detected as a positive crossing of the 0mV threshold. For each morphologically-detailed model, we manually defined n=6 dendritic branches that corresponded to a unique straightforward path starting near the soma (<30μm) and ending in a distal branch termination (>200μm). Branches were insured not to overlap with each other. Synapses were inserted on the morphology according to their locations on the dendritic tree on the Electron-Microscopy dataset. Per dendritic branch, a “uniform” surrogate of synaptic distribution was also created by randomly picking locations from a uniform distribution of distance to the soma along the branch (see inset of **Fig. 4A**). In **Fig. 4E-G** and **Fig. 5A-D**, synapses were randomly split into 80% excitatory and 20% inhibitory synapses.

Synaptic currents from AMPA, GABA and NMDA receptors were taken from (*61*) and inserted in the IN models (see properties in **Table S3**). For simplicity, NMDAR currents were only included in the SST-IN model and not in the PV-IN model (**Table S3**, see also **Fig. 1G**). For the SST-IN model in absence of NMDAR (condition “no NMDA, AMPA+”, see **Fig. 4E-G**), the AMPA weight was increased by 50% to match the peak amplitude of PSPs at -60mV.

Background activity in **Fig. 4E-G** was achieved by stimulating each synapse with an independent homogeneous Poisson point process. Rates of Poisson processes were adjusted to bring the neurons in the -60mV range. For the PV-IN model, this was achieved F_inh_=2.25mHz per synapses and F_exc_=3mHz per synapse. For the SST-IN model, this was achieved with F_inh_=8mHz per synapses and F_exc_=1mHz per synapse. In **Fig. 4F**, spiking response curves were fitted to the function:

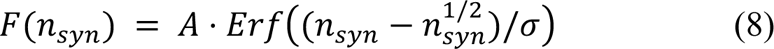

where *Erf* is the error function (fitting with the method=’*Nelder-Mead’*’ from the *scipy.optimize.minimize* function).

In **Fig. 5A-D**, the synapses were now driven by inhomogeneous Poisson processes, i.e. where they were controlled by time-varying “input” rates (shown in grey in **Fig. 5A,B**). In **Fig. 5A**, the excitatory and inhibitory rates are driven by step functions. In **Fig. 5B**, the rates are driven by an Ornstein-Uhlenbeck stochastic process (i.e. temporally-correlated Gaussian fluctuations) of time-constant τ=50ms with mean μ=2 and variance σ^2^=0.75^2^ multiplied by a scaling factor. This scaling factor for synaptic activity rates (in mHz/synapse, see scale bars in **Fig. 5B**) was adjusted so that the output firing rate reached 20±1Hz in all models considered (“PV”, “SST”, “SST:noNMDA, AMPA+”). The output rate function was then obtained by binning spikes from n=50 trials (“psth: per-stimulus time histogram”) and smoothing the resulting trace with a Gaussian filter of width 10ms. The cross-correlation functions were fitted to Lorentzian function 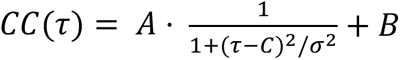 (constrained fitting with the method=’*L-BFGS-B*’ from the *scipy.optimize.minimize* function) where σ is the width parameter that provides an estimate of the temporal extent of the correlations to the input signal (**Fig. 5C**).

## Data analysis

### Synapse Distribution Analysis

We analyzed synaptic distributions on IN dendrites in a publicly-available 1mm^3^ reconstruction of the mouse visual cortex from a serial section transmission Electron Microscopy volume (*44*), i.e. a reconstruction with synaptic resolution. To specifically analyze INs, we also benefited from the manual proofread annotations of morphologically-identified INs found in (*45*) available at https://zenodo.org/record/7641781#collapseTwo. As a proxy for PV- and SST-INs, we considered the most numerous morphological type in each of those molecular classes, i.e. Basket and Martinotti cells respectively. From Ref. (*45*), n=59 Basket cells and n=41 Martinotti cells were available. For each cell, we benefited from a mesh reconstruction of its whole morphology as well as the set of synapses targeting this cell (“pre”) or targetted by this cell (“post”) with their localization in the volume (*45*). In **Fig. 3G,H**, we show such mesh reconstruction with the “pre” synapses (note the presence of both the axonal and dendritic trees). As a first step, we converted the mesh reconstructions of each cells into a “skeleton”, i.e. a set of segments with diameters, using the package MeshParty (https://meshparty.readthedocs.io/, version 1.16.4) as this enables to easily measure quantities such as “path length” along the dendritic tree (*44*). Next, we splitted axonal and somato-dendritic compartments based on the density of “pre” and “post” synapses per compartment (“pre” is high in dendrites/soma, “post” is high in axons) using the function *meshwork.algorithms.split_axon_by_annotation* from MeshParty. The few synapses (<1%) located on the axonal compartments were excluded from further analysis. We next focused only on the somato-dendritic compartments only. To analyze synapses along the dendritic path (**Fig. 3I,J**), the somato-dendritic skeleton of each cell was divided into a set of “dendritic cover paths” covering the full dendritic tree (without any overlap). On all segments of this unique “dendritic cover path”, we could compute the “path distance to soma” and count the number of synapses in the segment to report either the linear density of synapses (**Fig. 3I**) or the absolute count (**Fig. 3J**) as a function of the distance to soma.

### Visually-evoked spiking dynamics of PV- and SST-INs during natural movies

To analyze the temporal dynamics of the spiking activity in PV- and SST-INs, we re-analyzed the publicly-available *Visual Coding* dataset from Ref. (*50*) of Neuropixels recordings with visual stimulation and phototagging of INs. We considered the sessions in both the “brain_observatory1.1” and the “functional_connectivity” datasets performed on Pvalb-IRES-Cre;Ai32 (N=8 session) and Sst-IRES-Cre;Ai32 mice (N=12 sessions). We first restricted the analysis to spiking units in the visual cortex (i.e. annotated as belonging to one of the visual areas: “VISp”,“VISal”, “VISam”, “VISl”, “VISli”, “VISmma”, “VISmmp”, “VISp”, “VISpm”, “VISr”, see (*50*)). We then splitted single units into positive and negative units with respect to the molecular marker for INs. To this purpose, we analyzed the phototagging protocol (specifically the 10ms light pulse stimulation) and computed the firing in the pre-stimulus [-10,0]ms and post-stimulus [0,10]ms window. Units were defined as positive if their post-stimulus firing was above 20Hz and twice larger than the pre-stimulus firing (see “phototagging” insets in **Fig. 5F**). Using this criterion, we found n=146 PV-positive units over N=6 sessions (n=24±14 units/session, two sessions in Pvalb-IRES-Cre;Ai32 mice were excluded from the analysis because they had less than 2 positive units) and n=97 SST-positive units over N=12 sessions (n=8±6 units/session). To estimate local cortical activity, we considered a random subset of n=100 negative units in each session (out of the n=259±41 negative units/session in Pvalb-IRES-Cre;Ai32 mice and the n=308±108 negative units/session in Sst-IRES-Cre;Ai32 mice). We next analyzed the spiking dynamics during the two natural movies that were presented in all sessions: “natural_movie_one” and “natural_movie_three”. For each movie, we concatenated the spike times of the different units and different movie presentations to build an average time-varying rate for a specific IN population (see **Fig. 5F**). This time-varying rate was smoothed by a Gaussian kernel of width 10ms. We did this either by pooling all units of all sessions for a single stimulus (shown in **Fig. 5G**, resulting in a stimulus-evoked dynamics) or on a per-session basis (shown in **Fig. 5F**) to be able to analyze properties across sessions (**Fig. 5H**). As in the model (**Fig. 5C**), the single-session cross-correlation functions were fitted to a Lorentzian curve to estimate the temporal extent of correlations (**Fig. 5H**).

### Analysis of fluorescence signals

Two-photon imaging recordings of calcium activity were preprocessed with the suite2p software (*75*) to perform the registration and the extraction of calcium signals. Identification of SST+ cells were performed using the cellpose algorithm (*76*) on the time-averaged image of the field of view. Raw fluorescence traces were corrected for neuropil contamination by subtracting Suite2p-neuropil traces using a fixed scaling coefficient of 0.7 (*75*). To be able to compare activity across cells and mouse lines, fluorescence signals were normalized. To do this we used the ΔF/F_0_ method, calculated using the following formula: (F-F_0_)/F_0_, where F is the fluorescence and F_0_ is the time-varying baseline fluorescence evaluated as the lowest 5th percentile with a 2min sliding window. After neuropil subtraction, calcium signals were deconvolved using the deconvolution algorithm oasis from ref. (*77*) with settings from ref. (*75*) a (*76*) kernel time constant of 1.3s. Visually responsive cells were defined as cells displaying a significant increase (Anova test, p<0.05) over the n=10 stimulus repetitions between the intervals [-1,0]s and [1,2]s preceding and following stimulus presentation respectively.

### PSD-95 Venus puncta detection

PSD-95 puncta quantification was performed in dendrites simultaneously labelled with td-Tomato). Td-Tomato fluorescence allowed effective dendrite tracing in order to estimate PSD-95 puncta density relative to soma distance. For a individual dendritic branches, image acquisitions were performed proximally (less than 40um away from the soma) and distally (approximately 100um away from the soma). To avoid errors while defining distal and proximal dendritic segments, we only imaged dendrites that did not cross other dendritic branches in the fields of view. In addition, PSD95 quantification was performed in single images and not in z-stacks. Image acquisition was performed for both td-tomato (excitation peak at 555nm) and ATTO647N for mVenus (excitation peak at 647nm). Images were subsequently analyzed using Fiji software using the Analyze Particules plugin. PSD-95 mVenus fluorescence images were first filtered using Fast *Fourier transforms* using the Fiji FFT BandPass Filter function. Large structures (low frequency) were filtered down to 60 pixels, small structures (high frequency) were filtered up to 2 pixels. The images were thresholded using a minimum intensity threshold of the mean plus three times the standard deviation of the background. Puncta detection was conducted using the Analyze Particles plugin in Fiji, considering puncta with areas larger than 0.01 square micron. No maximal size limit was defined. Puncta density was calculated by dividing the number of puncta counts by the length of the analyzed dendritic segment. The length of the dendritic segment was defined on the td-tomato fluorescence channel.

### Statistics

We used Wilcoxon rank-sum tests for independent group comparisons, Wilcoxon signed-rank tests for paired tests and Student’s t-tests for a single group analysis. No statistical methods were used to pre-determine sample sizes, but our sample sizes were similar to those used in previous publications. Allocation into experimental groups was not randomized. Data collection and analysis were not performed blind to the experimental conditions.

